# A bacterial extracellular matrix protein forms a supramolecular metallogel

**DOI:** 10.64898/2026.03.09.710396

**Authors:** Mnar Ghrayeb, Aliza Ashman, Soham Mukhopadhyay, Amos Felig, Juergen Jopp, Yael Levi-Kalisman, Uri Raviv, Luca Bertinetti, Yael Politi, Vasily Zaburdaev, Sharon Ruthstein, Liraz Chai

## Abstract

The microbial extracellular matrix (ECM) is a complex network of self-secreted biopolymers uniting the cells in biofilms, providing them with structural integrity, and contributing to their elevated resistance to antibiotic treatments. Recently, there is a growing realization that a regulated, bidirectional cross-talk of bacteria and ECM confers biofilms with tissue-like traits, however, the mechanisms of spatio-temporal self-organisation of ECM and its regulation are still poorly understood. In the model organism for biofilm formation *Bacillus subtilis*, TasA is the major protein component of the extracellular matrix. We recently showed that TasA, isolated in the form of stable and structured globules, assembles into elongated and ordered fibers via a donor-strand complementation mechanism. In this study, we discovered that in the presence of zinc metal ions, TasA is able to form hydrogels with > 97% water content. Electron- and atomic force-microscopies as well as small angle X-ray scattering measurements show that cross-linking with zinc ions induces a transition in TasA morphology from one-dimensional fibers to two-dimensional sheets. Electron paramagnetic resonance measurements then show that such a significant morphological shift is associated with molecular changes in the coordination environment of zinc ions, which lead to structural changes at the protein level. When assembling into macroscopic networks, TasA-Zn metallogels exhibit viscoelastic properties and a fast recovery following an excessive strain. These metallogels represent a novel class of bacterially-derived ECMs that form easily at room temperature without covalent crosslinking, and may be used as a natural matrix-mimics in biofilm models for infection studies.

## Introduction

The extracellular matrix (ECM) is a three-dimensional network of biopolymers that surrounds cells in eukaryotic tissues and provides them with structural and mechanical robustness (1). Tissue ECMs are crucial for their homeostasis, morphogenesis and regeneration, as well as for cell differentiation, growth, and movement (2,3). This is also the case in the microbial world of bacterial biofilms, where cells are encased in a self-secreted ECM, commonly composed of proteins, polysaccharides, and nucleic acids (4,5). Concurent with the emerging perspective of biofilms bearing analogy to eukaryotic tissues, there is growing interest in better understanding the processes governing ECM assembly, self-organisation, and regulation (5, 6). However, while it is recognized that ECM plays a remarkably broad variety of roles, very basic processes related to its individual components are only being discovered in biofilms. (4, 6–10 TasA is the major ECM protein component in biofilms of the Gram-positive bacterium *Bacillus subtilis*, serving as a model organism for biofilm formation. Originally identified in *B. subtilis* spores (11), TasA also forms fibrils in the extracellular space of biofilms (7, 12, 13). Starting from observations of soluble oligomers of isolated TasA (14), later to be discovered as a jelly-roll of *β*-sheets (15), we have identified environmental conditions that lead to TasA aggregation *in vitro*, and showed that it is polymorphic in its aggregation pathways, structure, and architecture (16).

While forming fractal colloid-like aggregates at low pH (17), it assembles into highly ordered fibrils in neutral/alkaline pH and elevated protein concentrations that bundle at high salt concentrations. Regardless of the assembly pathway, TasA fibrils share a similar X-ray diffraction fingerprint (18), which we recently attributed to a donor-strand complementation mechanism (7). Interestingly, a similar fibrilation mechanism has recently been observed with a protein of archael origin (19, 20). Here we report on the discovery that, mediated by zinc ions, TasA has the capacity to form a hydrogel, thereby expanding its polymorphism repertoire from monomers, through oligomers, and fibrils, to (multi)fractal cross-linked networks. The involvement of zinc is not coincidental. We have previously detected zinc accumulation in whole biofilms (18, 21) while the structure of TasA fibers indicated putative zinc-binding sites (7). Here we used electron- and atomic force-microscopies together with X-ray scattering to show that, microscopically, the transition to the gel state is associated with a transition from fiber-dominated structures to two-dimensional sheets in a zinc-concentration dependent manner. Surprisingly, X-ray diffraction analysis suggests that such drastic macroscopic morphological changes involve only marginal structural changes of the TasA monomers at the molecular scale; these structural changes are enabled by a change in the coordination environment of zinc ions, as revealed by electron paramagnetic resonance measurements.

TasA-Zn metallogels form under normal room temperature conditions, convey material properties in a range comparable to that of common ECM-mimicking gels, and due to their metal-ion-dependent polymerization mechanism, they are self-healing: showing fast recovery following an excessive strain. These metallogels represent a novel class of bacterially-derived ECMs, which may also become a crucial advance in the development of 3D biofilm models (22) for studies of bacterial infections.

### The extracellular bacterial protein TasA aggregates upon the addition of zinc ions

Our previous studies indicated that TasA aggregation may be associated with zinc salt solutions. Specifically, we have shown that zinc is accumulated in biofilms (18,21), TasA is co-isolated with zinc (21), and finally, we have identified a putative zinc binding site in the cryo-EM structure of TasA fibrils (7). These findings were consistent with the homology of TasA to camelysin, a Zn-dependent metalloprotease (12,15). We therefore asked if the purified TasA aggregates upon the addition of zinc ions *in vitro* (see Fig. 1A for two steps of the aggregation protocol). We monitored the solutions of TasA following mixing with ZnCl_2_ solutions. Figures 1B,C show aggregates of TasA that form when mixing 20 mM ZnCl_2_ with 12 µM purified TasA.Fig. 1B shows an optical microscopy image of TasA aggregates that form immediately after mixing a TasA solution with ZnCl_2_ (step 1 in the schematic of Fig.1A). The aggregates were visible both in bright field and in fluorescence modes (the bright field and fluorescence channels are overlaid, Fig. 1B), where for contrast enhancement, Rhodamine B was added to a TasA solution prior to the zinc addition. TasA aggregates were merged into a semi-solid bulk by centrifugation (Fig. 1A, step 2 in the schematic). This semi-solid material resembled a gel according to a tube inversion test (Fig. 1C).

**Fig. 1.**
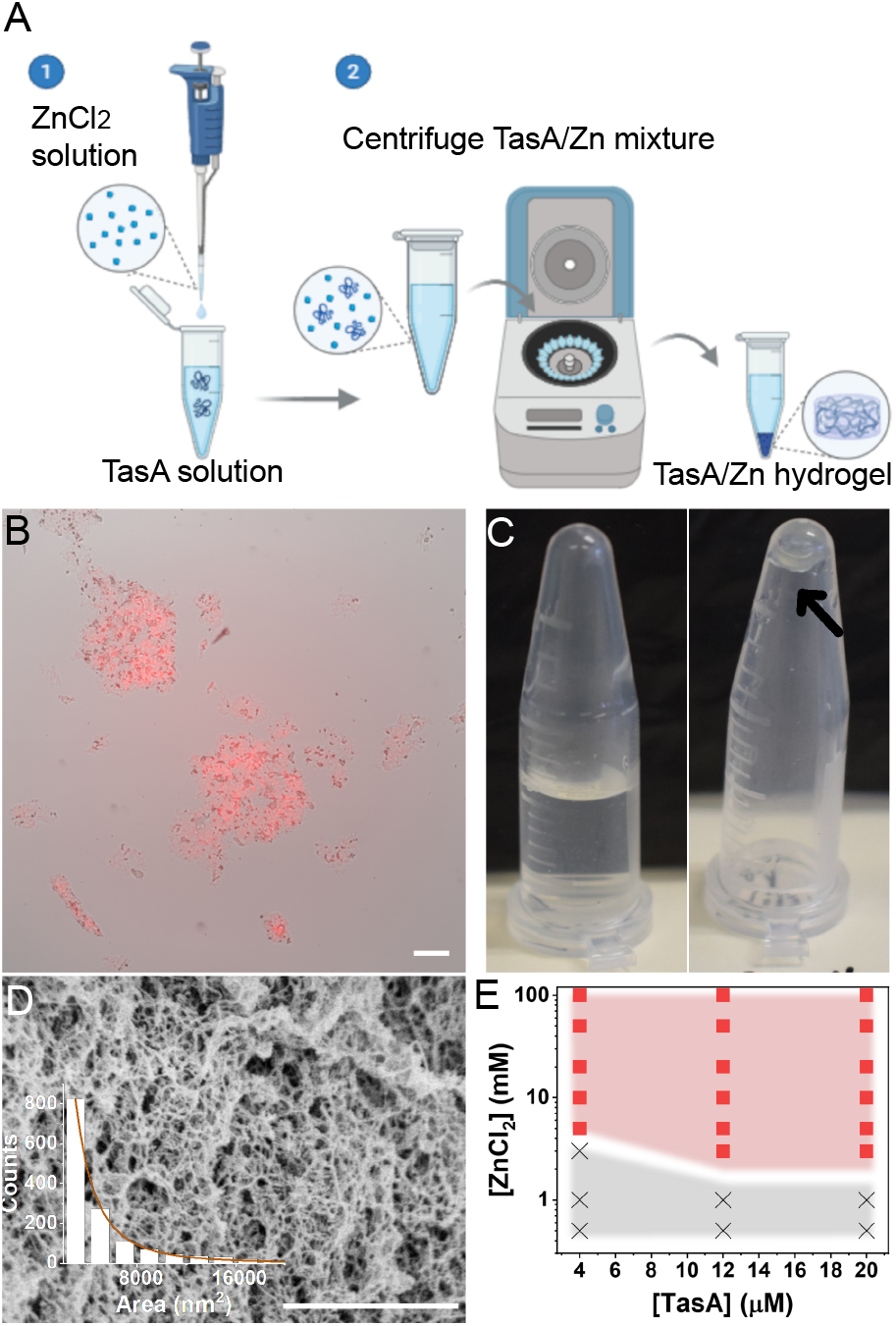
TasA aggregates in the presence of zinc chloride. Schematic illustration (A) of the steps used to induce TasA aggregation in solution (1) and combining the aggregates into a bulk (2). Bright field (B) image of aggregates forming as 20 mM ZnCl_2_ drops are added to 12 µM TasA stained with 10 µg/mL Rhodamine B, overlaid with fluorescence microscopy image of aggregates stained by Rhodamine B. Scale bar corresponds to 100 µm. (C) Inverted tube assay of 12 µM TasA with 1 mM ZnCl_2_ (left) and 20 mM ZnCl_2_ (right), the hydrogel is marked with an arrow. SEM image of a TasA hydrogel and analysis of the pore area in the gel shown in the insert. Scale bar corresponds to 1 µm (D). Summary of TasA gelation conditions (E), where formation of TasA-Zn aggregates is shown as a function of the tested concentrations of both TasA and ZnCl_2_. Red squares indicate conditions where aggregates formed, whereas gray crosse.s indicate no aggregation (Fig. 3).

Scanning Electron Microscopy (SEM) revealed that the TasA aggregates were composed of a mesh-like network, consistent with a characteristic gel morphology (Fig. 1D). We used SEM images of TasA networks to calculate their pore size distribution (see Fig. S1 for an example of the image analysis), revealing a mesh size ranging between ~ 20 nm - 500 nm (assuming circular pores), with the lower bound to the pore size being limited by the image resolution (See Fig. S2 and table S1 for SEM images and pore size distribution in TasA aggregates formed by mixing TasA and ZnCl_2_ at varying concentrations).

To account for the range of TasA and ZnCl_2_ concentration mixtures that lead to aggregation, we screened the concentrations of TasA and zinc chloride couples using optical density (OD) as a readout of aggregate formation. TasA (4-20 µM) and ZnCl_2_ (0.5-100 mM) solution mixtures were incubated without shaking at room temperature, and their OD was measured after 18 hours (see SI Table S2 for the OD values). We determined the cutoff for aggregation as OD > 0.12, (which is when aggregates became visible) and summarized the results in Fig.1E; the concentration couples of TasA and ZnCl_2_ that resulted with aggregation are marked by squares and by crosses otherwise.

This summary shows that in the tested range of TasA concentrations (4-20 µM), aggregates formed at a critical ZnCl_2_ concentration of ~3 mM, considerably smaller than the 0.5-1.5 M NaCl concentration, necessary to form TasA fibrils with similar TasA concentrations (7, 16) (3 mM ZnCl_2_ has a screening length equivalent to 12 mM NaCl). An additional indication that aggregation is concentration-dependent is the weight of TasA aggregates that for the tested concentrations (4 µM, 12 µM, 20 µM) increased with ZnCl_2_ concentrations (0.5-100 mM) until saturation at ~ 20 mM ZnCl_2_ (Fig. S3). Among other salts we tested, (CaCl_2_, MgCl2, MnCl_2_, FeCl_3_, FeSO_4_, NiCl_2_, and CuCl_2_), only copper ions were able to induce gelation, however, at higher threshold concentrations (>5 mM). We continued to focus on ZnCl_2_ for further analysis, and following the observation that TasA-Zn aggregates resembled hydrogels in morphology, we next tested the material properties of TasA-Zn aggregates.

### Characterization of mechanical properties and water content of TasA-Zn aggregates reveals their gel-like properties

We performed rheology measurements of TasA-ZnCl_2_ mixtures as a function of time to examine whether they exhibited a sol-gel transition dynamics (Fig. 2A). Specifically, we used 4 µM TasA, which is a low enough concentration that allowed us to resolve temporal changes of mechanical properties en route to gelation. We added ZnCl_2_ to different final concentrations in the range of 1-20 mM, and measured the temporal variation of the storage (*G*^*′*^) and loss (*G*^*′′*^) moduli, reflecting the elastic and viscous characteristics of the hydrogel, respectively. We note that there was a large variability in the results obtained for each TasA-ZnCl_2_ couple. This is due to the sample preparation corresponding to step 1 in Fig. 1 A. To capture the time lapse of the gelation process, here we worked with loose aggregates that formed in an epitube as well as on the rheometer while we probed them (as in Fig. 1B) and not with the centrifuged bulk gel samples (as in Fig. 1C) (see Methods).

**Fig. 2.**
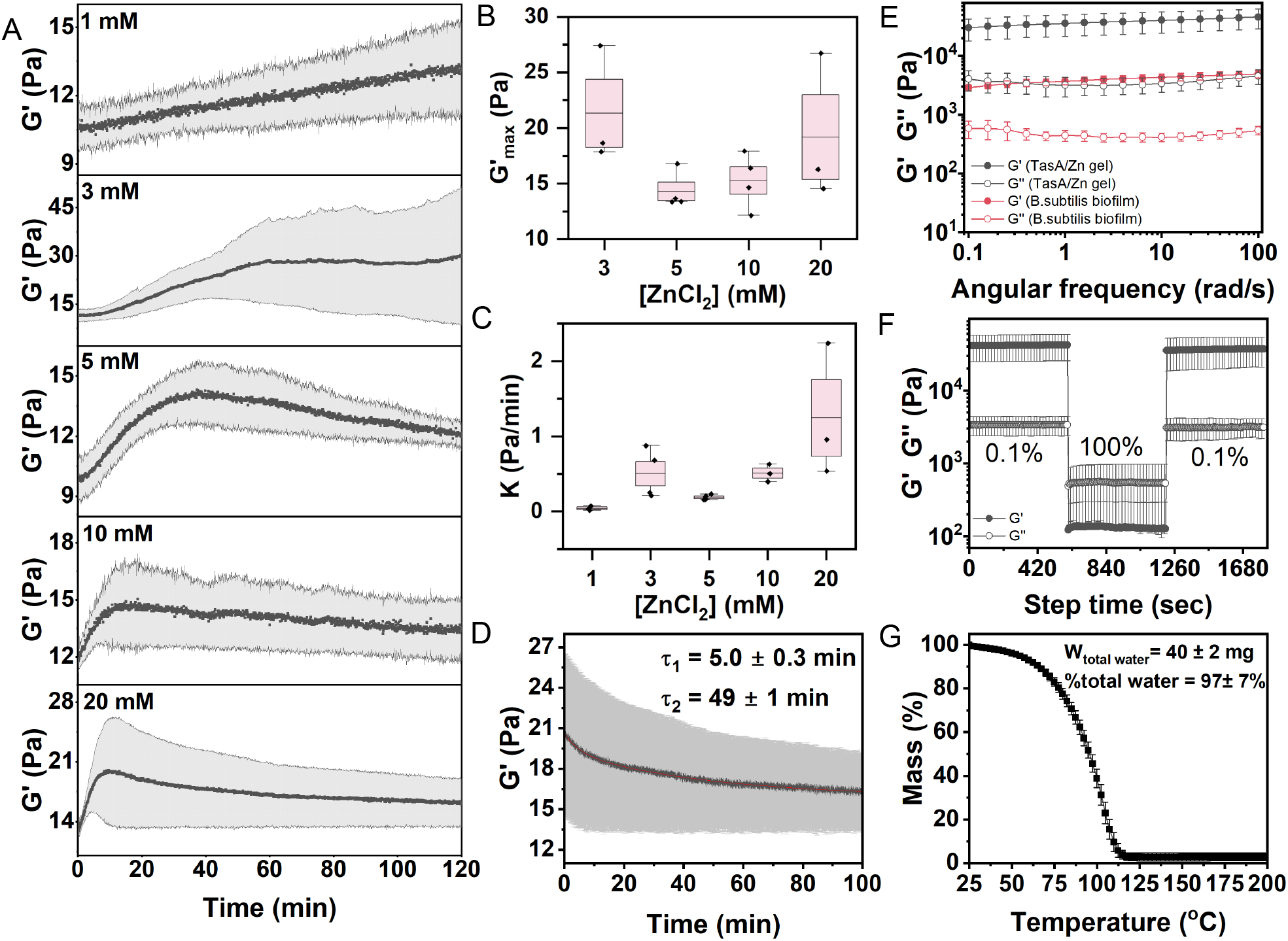
Characterization of TasA-Zn aggregates using rheology and thermogravimetric analysis reveals their hydrogel properties. Rheology measurements showing the storage modulus (*G*^*′*^) of a TasA-Zn gel formed with 4 µM TasA and increasing concentrations of ZnCl_2_ in the range 1 mM - 20 mM, plotted as a function of time. Average values are denoted by dark symbols, and the error is denoted by a gray area (A). Analysis of the *G*^*′*^ plots shown in A, highlighting the *G*^*′*^ maximal value (*G*^*′*^max) (B), *G*^*′*^ increase rate (*K*) as a function of ZnCl_2_ concentration (C) and the relaxation times obtained from a bi-exponential fit to the “tail” of *G*^*′*^(*t*) gelation curve for 12 µM TasA, 20 mM ZnCl_2_) (D). Rheology of a bulk gel, made from 12 µM TasA and 20 mM ZnCl_2_ (black symbols) together with rheology of wild-type (WT) *B. subtilis* biofilms (red symbols) (E). Recovery from damage of a TasA-Zn hydrogel tested by measuring *G*^*′*^ (full symbols) before, during, and after application of 100% strain (F). ThermoGravimetric Analysis (TGA) of a TasA-Zn hydrogel (G).

The general dynamics of *G*^*′*^ versus time (Fig. 2 A) was similar in all tested TasA-ZnCl_2_ couples, exhibiting a rise of *G*^*′*^ to a maximal value, which then decayed with time (see *G*^*′′*^ measurements in Fig. S4). The aggregation dynamics of TasA in the presence of zinc ions resembled that of gelation propagation of folded globular proteins as well as of ioniccross-linked hydrogels (23,24). Starting from a lag phase, attributed to diffusion limitation of the cross linker, there was an increase in the modulus (gel cross-linking) that peaked at the gelation point and then decayed as the network rearranged and cross linkers could be leaching or substituting. We barely observed a lag phase prior to the increase of *G*^*′*^, for example when 4 µM TasA was mixed with 3 mM ZnCl_2_, which indicates fast diffusion of ions and cross-linking. Interestingly, relaxation of *G*^*′*^ in gelled networks is a property of reversible cross linking, whereas in non-reversible, for example covalent-cross linking, *G*^*′*^ changes with time in a sigmoidal manner where *G*^*′*^ plateaus at a maximum value rather than relaxes (25): relaxation is highly unlikely to occur in covalently cross-linked hydrogels due to essentially permanent links and small chain lengths between them. In fact, reversible cross-linking in biofilm extracellular matrices would provide these networks with the necessary flexibility to form, break, and then reform, depending on environmental conditions.

Quantitative analysis of the gelation formation curves in Fig.2A (see Fig. S5) shows the variation of *G*^*′*^*max* (maximum value of *G*^*′*^) and *K* (defined as *G*^*′*^ increase rate) values as a function of ZnCl_2_ concentration (Figs. 2B, 2C, respectively). Statistical analysis of the data suggests that only the differences between *K* values are significant, indicating that the gelation rate zinc-ion concentration dependent, as expected in an ionic cross-linking gelation process. Interestingly, the network relaxation (we used only the fully cross-linked hydrogel with (4 µM TasA and 20 mM zinc chloride for this analysis) had two characteristic time scales, ~5 and ~50 minutes (Fig. 2D), which may be attributed to zinc substitution between two different zinc binding environments (see below).

We next characterized the mechanical properties of the bulk TasA-Zn hydrogel (obtained after centrifugation, step 2 in Fig. 1A) which, according to Figs.1E and S3, is fully cross-linked (formed by mixing 12 µM TasA, 20 mM ZnCl_2_). Frequency sweep measurements (performed at 0.1% strain Fig.2E, see Fig. S6 for the strain sweep measurement) show a typical viscoelastic solid-like behavior of the TasA-Zn gel with a larger elastic contribution (*G*^*′*^ > *G*^*′′*^) over a frequency range of 0.1-100 Hz. This, together with the parallel lines of *G*^*′*^ and *G*^*′′*^ suggest that the material is gel-like (26).

Interestingly, whole wild type (WT) *B. subtilis* biofilms also exhibited a viscoelastic gel-like behavior, with G’ = 2700 *±* 500 Pa and G” = 290 *±* 25 Pa, corresponding to the elastic and viscous moduli of TasA-Zn hydrogels formed by (4 µM< *T asA* <12 µM) and 20 mM ZnCl_2_ (fully-cross-linked)(see Fig. S6).

A self-healing test showed that TasA-Zn hydrogels recover immediately from a high (100%) shear strain (Fig. 2F), indicating that cross-linking is dynamic, supporting the non-covalent, zinc ion-mediated cross-linking mechanism, where the zinc ions quickly rearrange after shear and restore the original mechanical properties of the gel. This is an important trait for biofilms that reside in the soil and need to endure harsh shear rates. Indeed, whole biofilms also recover following a similar shear strain (Fig. S6).

An additional central property shared by both biofilms and hydrogels is their high water content. Therefore, we measured the water content of the TasA-Zn gels (12 µM TasA with 20 mM ZnCl_2_) using ThermoGravimetric Analysis (TGA). Fig. 2G shows the TGA curve and the calculation of water content, which was 97% *±* 7%, thus putting it in the range of high water-content gels. We henceforth refer to the material formed by centrifugation of the TasA and ZnCl_2_ aggregates as TasA-Zn gel.

To summarize, we have shown that addition of zinc ions to the solution of TasA protein in a broad range of concentrations results in sol-gel transition.The resulting metallogels are viscoelastic, self-healing, have a high water content, and are similar in their respective material properties to whole biofilms.

### From 1D fibrils to 2D sheets that assemble into 3D networks

Following the characterization of TasA-Zn as a bulk metallogel, we were intrigued to decipher the mechanism of aggregation of TasA into a hydrogel. We therefore visualized TasA using Cryo Transmission Electron Microscopy (TEM) in the absence and presence of ZnCl_2_ in varying concentrations. Figure 3 shows that TasA (12 µM) forms fibrils prior to ZnCl_2_ addition (Fig. 3A), in agreement with our previous observations (16), but when ZnCl_2_ is then added (1 mM), there is a coexistence of TasA fibrils, clusters, and occasionally bundles (Fig. 3B, see also Fig. S7). As the concentration of ZnCl_2_ is further increased (20 mM), the most dominant morphology resembles thin crumpled sheets (Fig. 3C), which is radically different from fibrils (Fig. 3A). At these TasA (12 µM) and ZnCl_2_ (20 mM) concentrations, TasA is already in the form of a gel (Fig. 1E), indicating that the transition from TasA in solution to TasA gels involves a transition from fibrils to two-dimensional sheets. These then further assemble into higher hierarchies to form a gel-like network (see e.g., SEM image, Figs. 1D, S2).

**Fig. 3.**
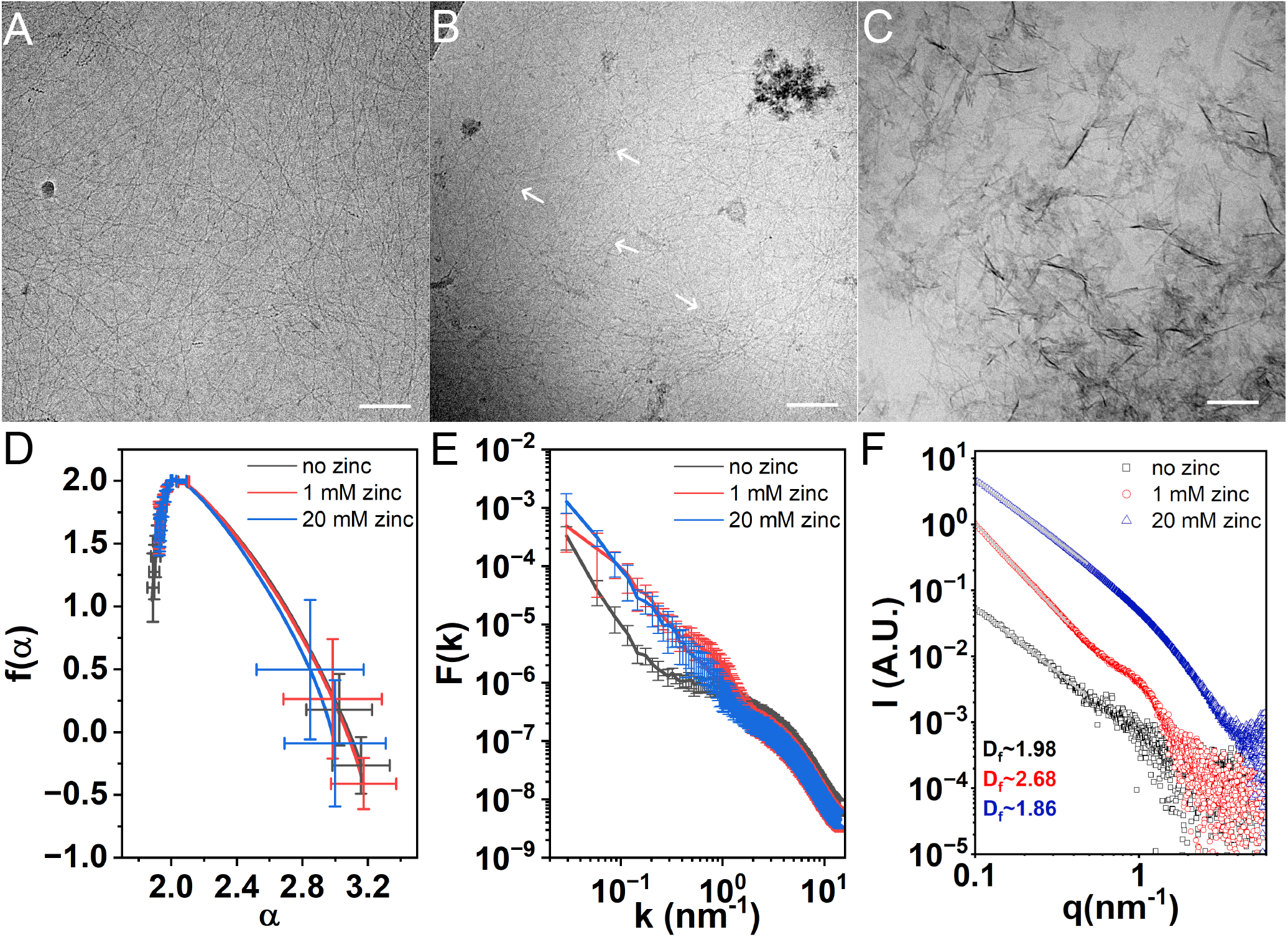
Cryogenic transmission electron microscopy (Cryo-TEM) and small-angle X-ray scattering (SAXS) analysis of TasA in the presence of zinc chloride solutions. Cryo-TEM images of 12 µM TasA with 0 mM (A), 1 mM (B), and 100 mM (C) ZnCl_2_, showing the transition of TasA from fibers (A) to a mixture of fibers and clusters (B) and to 2D sheet-like structures (C), scale bar corresponds to 100 nm. Multifractal spectrum *f* (*α*) as function of fractal dimension exponent *α* (D) and structure factor as a function of the absolute value of wave-vector *k* (E) of TEM images (see also Fig. S8). (F) Azimuthally integrated, background-subtracted SAXS intensity curves as a function of the magnitude *q* of the scattering vectors from 12 µM TasA with the indicated ZnCl_2_ concentrations (N = 9 ROIs for D and E, and N = 20 for F). The slopes of the linear fit to the low *q* scattering data (solid lines) provide the fractal dimension of the samples (*D*_*f*_).

TEM images of clusters for varying ZnCl_2_ concentrations show complex hierarchical structures which could be a signature of a fractal or even a multi-fractal organisation. Recently, we have shown that organisation of complex biopolymers or microphase separating systems, such as chromatin in the cell nucleus, is consistent with a multi-fractal behavior (27). In a multifractal object, a whole spectrum of fractal exponents is required to capture the complexity of its structure across scales and its subregions (28). Indeed, the quantitative analysis of TEM images reveals a multifractal behavior both in conditions without the addition of zinc ions as well as in the metallogel regime, where we focused the analysis on the prominent clusters. Figure 3D (see the library of analyzed ROIs in Fig. S8) shows the multifractal spectrum *f* (*α*) where *f* is the fractal dimension of a subset of the image where the density exhibits local scaling with a singularity exponent *α* as a function of the observation scale (see (28) and SI for methodology details). While for monofractal or non-fractal structures the spectrum would converge to a fixed value, multifractals present a broad distribution as seen in Fig.3D.

A complementary way to quantify the structure of the aggregates is by the structure factor *F* (*k*) (which is calculated via the Fourier transform of the density-density correlation function), Fig. 3E. Consistent with the multifractal statistics in these images, there is no distinct power-law behavior at low values of the wave vector *k* which would be apparent in a monofractal case. All conditions share a similar hump at the tail of distributions for large *k* ≳ 2nm^*−*1^). Translated in the space scale of ≲ 3 nm it roughly corresponds to the size of TasA monomers. *F* (*k*) of no Zn condition lacks signal in low *k* values (*k* ≲ 1nm^*−*1^) as compared to metallogel conditions which exhibit large scale clusters. Interestingly, no Zn and 1 mM ZnCl_2_ had an additional “hump” in 0.4-2 nm^*−*1^ range whose location could be related to the TasA monomers and their bundles with diameter of about 3-16 nm) (7, 12, 16), which disappeared at 20 mM ZnCl_2_, as fibers were largely replaced by sheets.

Whereas TEM image analysis already provides interesting hints to the aggregates’ structure across scales, it is still effectively a two-dimensional projection of the actual three-dimensional mass density distribution. In particular, it makes the interpretation of the multifractal spectrum challenging. To directly access the 3-dimensional structure of the TasA gels, we measured the solution small-angle X-ray scattering (SAXS) curves at the same concentrations used for TEM imaging (12 µM TasA, and 0 mM, 1 mM, 20 mM ZnCl_2_) (Fig. 3F). First, we observed that at low scattering vectors *q* the SAXS intensity well-fitted a power-law. The slope of a linear fit to the SAXS data (in a double logarithmic scale) provided the mass fractal dimension *D*_f_ of the sample (29), associated with the 3D arrangement of the protein. Although generally of the order of 2 (consistent with the position of observed maximum in the singularity spectrum around *α~* 2), values of *D*_f_ are distinct and, supported by the TEM images, they provide hints towards the mechanism of TasA assembly. In the lack of ZnCl_2_, the SAXS-based, *D*_f_ = 1.98, suggests that TasA fibrils (observed in Fig. 1A) were densely packed to the level that they appear as a continuous 2D mesh. As the ZnCl_2_ concentration was increased to 1 mM, the *D*_f_ = 2.66 in solution was consistent with the dominance of 3D clusters in solution, observed by TEM (Fig. 3B, and Fig. S7). The fractal dimension following a further increase in ZnCl_2_ concentration (20 mM) was *D*_f_ = 1.86, indicating a two-dimensional morphology, which, unlike the arrangement of monomers (when no zinc chloride was added), originated from the formation of 2D sheets, clearly observed by TEM (Fig. 3C).

Consistent with this explanation and the structure factor analysis of the TEM images, we also observed a clear “hump” in the SAXS data at intermediate *q* values, corresponding to the size of TasA monomers, which disappeared at higher zinc ion concentrations.

These results indicate that the range of transitions from monomers to filaments, to fibers (bundles of filaments), and then to two-dimensional sheets eventually assembling in (multi)fractal hierarchical structure in the metallogel are driven by the changes in protein self-interactions mediated by zinc ions. This led us to investigate the role of zinc ions in shaping TasA structure at the molecular scale.

### Molecular origins of TasA-Zn metallogel formation

The significant morphology changes of TasA from fibrils to sheets with increasing zinc ions concentrations, prompted us to test whether zinc binding to TasA is coordinative and specific to certain functional groups in the protein sequence or whether it is a non-specific coulombic interaction. In proteins, zinc ions are typically coordinated by ligating residues, most commonly cysteine (Cys, via sulfur), histidine (His, via nitrogen), aspartate (Asp, via oxygen), and glutamate (Glu, via oxygen) Lacking cysteine residues in TasA marks His, Asp and Glu as potential zinc ions binding candidates with either His binding occurring through a nitrogen atom or via oxygen atoms in carboxylic acid groups of Asp and Glu residues.

To probe the coordination environment of metal ions in TasA gels and fibers, we conducted low-temperature electron paramagnetic resonance (EPR) spectroscopy using paramagnetically active Cu^2+^, which is commonly used as a substitute for diamagnetic and EPR-silent Zn^2+^ (30,31). Cu^2+^ was added to TasA being either in a fiber (no zinc ions added, 8 µM TasA) or in a gel state (following the addition of 20 mM zinc ions). Importantly, we did confirm that the addition of Cu^2+^ to TasA, similarly to Zn^2+^, induced fiber and gel formation (see Fig. S9). Figure 4A-B presents the EPR spectra of Cu^2+^ in Tris buffer for TasA fibers and TasA gels. By fitting spectral simulations to the data (32) we extracted g-values (see table. S3) and hyperfine coupling constants that reflect the nuclear environment coordinating the Cu^2+^ center (33).

**Fig. 4.**
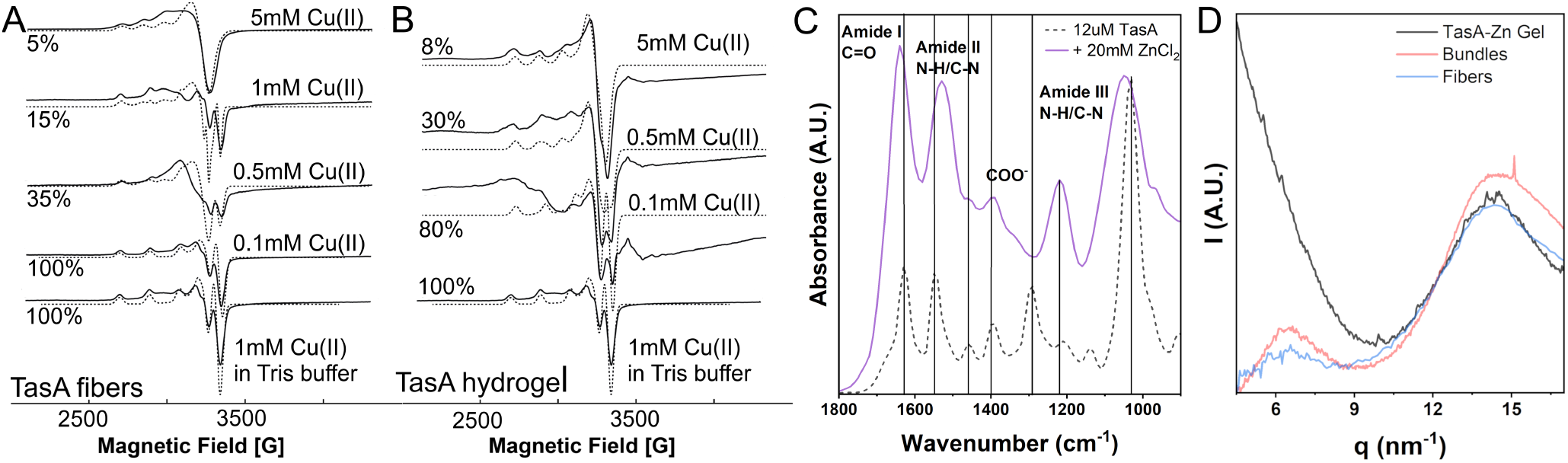
Molecular view into the interaction of TasA and Zn^2+^. Electron paramagnetic resonance (EPR) spectra of TasA in the form of fibrils following the addition of increasing CuCl_2_ concentrations (A) as well as of a TasA-Zn hydrogel that fromed with the addition of a mixture of a fixed ZnCl_2_ concentration (20mM) and increasing CuCl_2_ concentrations (B). The percentages near the plots represent the mobile copper ions in the Tris buffer, whereas the rest of the copper ions are bound to the protein in fibers (A) or gel (B) form. Fourier-Transform InfraRed (FTIR) spectra of TasA in the lack and presence of zinc ions (D). Dominant FTIR shifts: amide I stretching peak position (amide I) from ~ 1630 cm^*−*1^ to ~ 1650 cm^*−*1^; reduction and broadening of the COO^*−*^ symmetric stretch (~ 1390 cm^*−*1^, attributed to Asp/Glu; the N–H bending and C-N stretching modes shifting from 1550 cm^*−*1^ to 1520 cm^*−*1^ (amide II) and from ~ 1220 cm^*−*1^ to ~ 1250 cm^*−*1^ (amide III). SAXS/WAXS of TasA in the form of fibers and bundles versus TasA-Zn gel (E).

For TasA gels and fibers, EPR spectra were fitted using a two-component model: free Cu^2+^ in buffer and Cu^2+^ bound to TasA. The simulations were best fitted with axial symmetry, indicating octahedral coordination of Cu^2+^. Increasing Cu^2+^ concentrations led to greater protein coordination in both fibers and gel states, as indicated by the reduction in the fraction of free Cu^2+^ and corroborating its incorporation into fibers and gels. The extracted parameters suggest a coordination involving 1N3O or 2 N2O in fibers, in contrast to 2N2O or 3N1O in gels. The presence of an additional nitrogen in the direct coordination environment in gel versus fiber conditions, may suggest protein crosslinking with adjacent monomers contributing their His or backbone nitrogen to the zinc coordination pocket. Notably, the EPR spectrum at 5 mM Cu^2+^ in fibers is broader than in gels, implying possible exchange interactions between proximate copper ions in mobile fibers. In contrast, capturing Cu^2+^ when TasA polymers are crosslinked in a gel limits their mobility.

Complementary evidence for zinc coordination by TasA is provided by X-ray Photoelectron Spectroscopy (XPS) and Fourier-Transform InfraRed (FTIR) spectroscopy. XPS shows that transition of TasA from fibrilar to hydrogel form was associated with N 1s peaks (attributed to His) shifting to higher binding energy, and C 1s peak (carbon in Asp/Glu carboxylic acids) shifting to lower binding energies (Fig. S10 and table S4 with functional group assignment to XPS peaks (34)). The O 1s peak split shift is indicative either of zinc ions binding by Asp/Glu in a monodentate fashion (35, 36) or of zinc ions binding via Asp/Glu as well as via oxygen in water molecules (37).

FTIR spectra of TasA fibers before (black) and after (purple) the addition of zinc ions show clear shifts in the amide (I, II, III) carbonyl modes, which could be attributed to backbone carbonyl, as well as in the the N–H and C-N modes (Fig. 4C), typically attributed to Asp/Glu zinc binding (38) (see Fig. S11 showing that this effect is zinc-concentration dependent), thus marking Asp/Glu as possible ligands for zinc ions.

Altogether, EPR, XPS and FTIR data imply that zinc ion coordination occurs through backbone or Asp/Glu carboxylate groups and/or via His- nitrogen. Given our results and leaning on literature knowledge of zinc ions often coordinated in a 3N1O octahedral configuration in metalloproteinases (39, 40), a similar zinc binding site in TasA that would support cross-linking involves three equatorial His-nitrogen atoms and one oxygen atom originating from Asp/Glu in TasA, as well as two axial water molecules. A single TasA monomer has only two His residues, which would require an additional His nitrogen from a neighboring TasA monomer to complete the octahedral zinc coordination, and account for cross-linking.

The changes in coordination environment of Zn ions as seen by EPR, XPS and FTIR between fibers and gels led us to hypothesize that TasA may need to undergo a structural change to accommodate these atom rearrangements and facilitate formation of sheets. To this end, we used X-ray diffraction (XRD) to probe the structural changes within TasA in the states of fibrils and hydrogels. Fig. 4D shows the SAXS/WAXS of TasA-Zn gel (black) together with TasA fibrils (18) (no added zinc) (red, blue color) for comparison. We have previously attributed the two XRD reflections measured with TasA fibrils, at positions 6 nm^*−*1^ and 13.8 nm^*−*1^, to the inter-(beta) sheet and inter-strand (intra-sheet) separations in the repeating fibril subunit, respectively (18). Strikingly, while the XRD signal of a TasA-Zn gel retains the 13.8 nm^*−*1^ reflection, it lacks the 6 nm^*−*1^ reflection, as determined by peak deconvolution (Fig. S12). Furthermore, comparison of the XRD signals of TasA-Zn gel and WT biofilms (Fig. S12) showed a similar peak-loss in some areas in 12 h old biofilms. The disappearance of this peak, which we previously related with the lack of TasA expression, may now be explained by the presence of TasA in a different polymorph.

## Discussion

TasA is a *B. subtilis* protein which was originally discovered in the spore coat (11) and then observed in the form of fibers in biofilms’ extracellular matrix (12,13). *In vitro*, it adopts different polymorphs, globular monomers or oligomers (14), as well as fibrils (7,16), depending on environmental conditions. The stability of TasA as a globular protein is indicative of an energy barrier that “traps” TasA in a monomer or oligomer form, which is overcome when the environmental conditions change. As the protein concentration increases and/or when salt (high NaCl concentration or low ZnCl_2_ concentration) is added to the solution (16,18), the monomers get closer together, allowing them to interact through a donor-strand complementation mechanism(7). Here we show that zinc ions drive TasA aggregation into fibrils at much smaller concentrations compared to sodium ions, however, this cannot be explained by the shorter electrostatic screening length of divalent versus monovalent ions. An explanation, supported by the fact that divalent ions other than zinc and copper do not induce aggregation, is that the interaction of TasA and zinc is specific to certain residues on the TasA sequence. We showed that upon interaction with zinc ions, TasA forms networks with gel-like properties, and, based on a broad range of evidence accumulated by complementary techniques, we would like to suggest a potential mechanism for the transition of TasA from fibers to gels.

TasA fibers resemble beads on a string (both by TEM and AFM, (7, 14, 16)), where TasA monomers (‘beads’) are bound together via a donor-strand complementation mechanism(7). When zinc ions are added to TasA solutions, TasA forms sheets as is clearly observed in the TEM images (Fig. 3).

This extreme change of morphology involves molecular changes to accommodate the transition from fibers to sheets, as is indicated by recruiting an additional nitrogen (EPR) to allow zinc ion binding as well as changes in carboxylic acid binding (XPS), all of them attributed to histidine, aspartic and glutamic acids, which are known to act as zinc ion chelators. These molecular changes may be essentially held responsible for cross-linking and pushing the preferred TasA assembly from 1D fibers to 2D sheets. Such a transition may be explained by the presence of several Zn^2+^ binding sites with different binding coefficients within TasA monomers. In such a scenario, fibrils form first upon zinc ions addition, via the binding sites with the higher coefficient, and when the Zn^2+^ concentration exceeds a threshold value, the binding sites of a lower coefficient then act to make the arrangement of TasA monomers into sheets more favorable. Such a mechanism where TasA harbors Zn^2+^ (un)binding sites of different coefficients is consistent with the different relaxation times post the gelation propagation (Fig. 4D).

The extreme change in morphology (from 1D fibers to 2D sheets) and the fact that the TasA sheets lose a clear monomer/bead structural signature (XRD) suggest that at high enough Zn^2+^ concentration TasA changes conformation to accommodate further zinc ions by exposing new binding sites and recruiting additional monomers to coordinate zinc ions, thereby achieving cross-linking.

Corroboration of this hypothesis comes from the interpretation of the XRD data. The characteristic separation that we attributed to the inter-sheet inside a TasA monomer is lost, suggesting that the jelly-roll structure of TasA monomers opens up en route to forming sheets and that zinc-mediated cross-linking then occurs via flattened TasA monomers, enabling the thin sheet morphology. The latter is clearly observed by TEM, but AFM imaging provides even a better visual image of the gelation process: fibers mixed with oligomers/monomers (Fig. 5A) become thinner with the addition of zinc ions, and the oligomers/monomers in the background seem to disappear (Fig. 5B), possibly as they are recruited into TasA aggregates due to better binding to zinc ions; the bead morphology of the fibers is lost as sheets are starting to appear (Fig.5C).

**Fig. 5.**
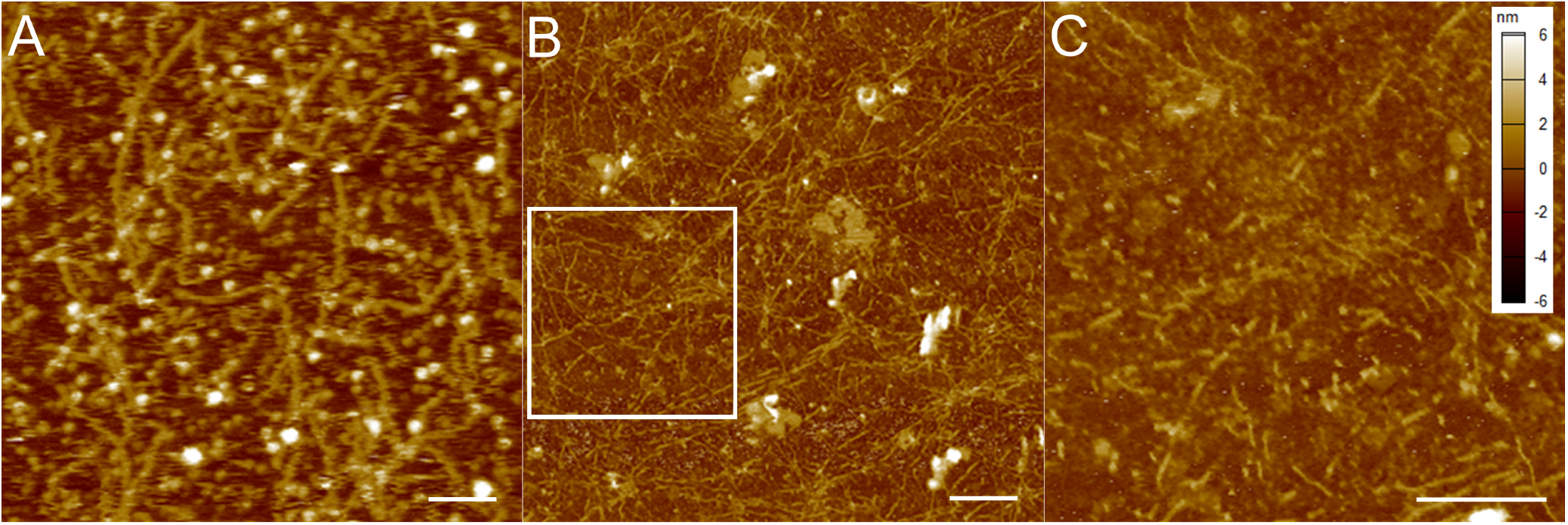
AFM imaging of TasA in the forms of fibrils and sheets. AFM images of TasA on mica, adsorbed from 6 µM solution (A), and following the addition of Zn^2+^(B); A zoom into the rectangle area in (B) is shown in (C), highlighting the morphology changes of TasA from 1D fibrils to 2D sheets. Scale bar corresponds to 100 nm.

Interestingly, it has been shown that archaeal extracellular proteins that assemble into fibrils via a donor-strand complementation mechanism, similarly to TasA, can assemble into higher hierarchical structures when exposed to Ca^2+^ ions, including folding into tubes(19) or networks(20). This strengthens the possibility that sheet formation via metal ion complex formation is not specific to TasA and that it may rather be shared by other extracellular fibril forming proteins.

This work is expected to contribute to both the biomaterials and biofilms fields. As a biomaterial, TasA-Zn metallogels expand currently used animal-, plant- and with very rare examples (20) of bacterial protein-based hydrogels to bacterially-derived *metallogels*; the latter being more relevant for bacterial cell growth in biofilm models, not only due to its bacterial origins but also because it does not involve neither chemical modification nor chemical cross-linking, it is water-based, and it forms easily at room temperature.

In the biofilms context, this study shows that TasA, hitherto known to assemble only into fibrils, also carries the capacity to form metallogels. While it is yet to be deciphered whether TasA-Zn metallogels are part of WT biofilm’s matrix, our findings exemplify for the first time that bacterial ECM protein components may assemble into supramolecular and reversible hydrogels, a trait which is highly useful for a living material that, whether in the ECM or as part of the spore coat, needs to assemble and disassemble according to rapidly changing environmental conditions.

## Materials and Methods

TasA was purified as described elsewhere (12,14) with modifications, as specified in the SI. The gelation protocol of TasA with *ZnCl*_2_ solutions was described in the text and it is further explained in the SI. Hydrogels were characterized using Scanning and transmission and Electron microscopies, rheology, Differential Scanning Calorimetry, Atomic Force microscopy, Electron Paramagnetic Resonance, and X-ray photoelectron Spectroscopy. See SI methods for manufacturers’ names. X-ray diffraction (XRD) measurements were performed at the ID13 beamline of the European Synchrotron Radiation Facility (ESRF) (Grenoble). SAXS data were measured in a flow-through capillary setup at the ID02 beamline (headed by T. Narayanan) at the ESRF (Grenoble, France) (41,42). The scattering intensity was azimuthally averaged and plotted as a function of the magnitude, *q*, of the scattering vector, 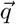. Before and after each measurement, the buffer of each solution was measured and subtracted from the measurement of the solution, as explained (43–46).

Detailed information on the experimental methods and sample preparation can be found in the SI Appendix.

## ACKNOWLEDGEMENTS

This work was supported by the Israel Science Foundation (grant 2090/24) to LC, the Deutsche Forschungsgemeinschaft (DFG, German Research Foundation) Grant 504222949 to LC, VZ, and MG in the framework of the Priority Program SPP2389 “Emergent functions of bacterial multicellularity”. MG acknowledges the support of the Neubauer Family Foundation for the PhD fellowship. We thank the European Synchrotron Radiation Facility (ESRF) for providing synchrotron radiation facilities, and we would like to thank Vitaly Gutkin for his help with XPS measurements and analysis, T. Narayanan and his team for assistance in using beamline ID02, May Yam Moshkovitz for her help with critical point drying of hydrogel samples, and Moshe Gottlieb for valuable discussions.

## References

1. Christian Frantz, Kathleen M. Stewart, and Valerie M. Weaver. The extracellular matrix at a glance. Journal of Cell Science, 123:4195–4200, 12 2010. ISSN 1477-9137. doi: 10.1242/jcs.023820.

2. Achilleas D. Theocharis, Spyros S. Skandalis, Chrysostomi Gialeli, and Nikos K. Karamanos. Extracellular matrix structure. Advanced Drug Delivery Reviews, 97:4–27, 2 2016. ISSN 0169409X. doi: 10.1016/j.addr.2015.11.001.

3. Fred T Bosman and Ivan Stamenkovic. Functional structure and composition of the extracellular matrix. The Journal of Pathology, 200:423–428, 7 2003. ISSN 0022-3417. doi: 10.1002/path.1437.

4. Anna Dragoš and Ákos T. Kovács. The peculiar functions of the bacterial extracellular matrix. Trends in Microbiology, 25:257–266, 4 2017. ISSN 0966842X. doi: 10.1016/j.tim.2016.12.010.

5. Hans-Curt Flemming, Eric D. van Hullebusch, Thomas R. Neu, Per H. Nielsen, Thomas Seviour, Paul Stoodley, Jost Wingender, and Stefan Wuertz. The biofilm matrix: multitasking in a shared space. Nature Reviews Microbiology, 21:70–86, 2 2023. ISSN 1740-1526. doi: 10.1038/s41579-022-00791-0.

6. Liraz Chai, Vasily Zaburdaev, and Roberto Kolter. How bacteria actively use passive physics to make biofilms. Proceedings of the National Academy of Sciences, 121(40):e2403842121, 2024. doi: 10.1073/pnas.2403842121.

7. Jan Böhning, Mnar Ghrayeb, Conrado Pedebos, Daniel K. Abbas, Syma Khalid, Liraz Chai, and Tanmay A. M. Bharat. Donor-strand exchange drives assembly of the tasa scaffold in bacillus subtilis biofilms. Nature Communications, 13:7082, 11 2022. ISSN 2041-1723. doi: 10.1038/s41467-022-34700-z.

8. Iztok Dogsa, Barbara Bellich, Mojca Blaznik, Cristina Lagatolla, Neil Ravenscroft, Roberto Rizzo, David Stopar, and Paola Cescutti. Bacillus subtilis epsa-o: A novel exopolysaccharide structure acting as an efficient adhesive in biofilms. npj Biofilms and Microbiomes, 10: 98, 10 2024. ISSN 2055-5008. doi: 10.1038/s41522-024-00555-z.

9. Ayantika Saha, Joshua M. Jones, Abigail Plummer, and Joseph W. Larkin. Formation of a swelling gel underlies a morphological transition in Bacillus subtilis biofilms. 2 2026. doi: 10.64898/2026.02.20.707077.

10. Macarena Siri, Agustín Mangiarotti, Anne Seewald, Nikolai Rosenthal, Shahrouz Amini, Emeline Raguin, Peter Fratzl, and Cécile M. Bidan. E. coli extracellular matrix: a tunable composite with hierarchical structure. 2 2026. doi: 10.64898/2026.02.22.707275.

11. Axel G. Stöver and Adam Driks. Secretion, localization, and antibacterial activity of tasa, a Bacillus subtilis spore-associated protein. Journal of Bacteriology, 181:1664–1672, 3 1999. ISSN 0021-9193. doi: 10.1128/JB.181.5.1664-1672.1999.

12. Diego Romero, Claudio Aguilar, Richard Losick, and Roberto Kolter. Amyloid fibers provide structural integrity to Bacillus subtilis biofilms. Proceedings of the National Academy of Sciences, 107:2230–2234, 2 2010. ISSN 0027-8424. doi: 10.1073/pnas.0910560107.

13. Natalie C. Bamford, Ryan J. Morris, Alan Prescott, Paul Murphy, Elliot Erskine, Cait E. MacPhee, and Nicola R. Stanley-Wall. <scp>tasa</scp> fibre interactions are necessary for Bacillus subtilis biofilm structure. Molecular Microbiology, 122:598–609, 10 2024. ISSN 0950-382X. doi: 10.1111/mmi.15315.

14. Liraz Chai, Diego Romero, Can Kayatekin, Barak Akabayov, Hera Vlamakis, Richard Losick, and Roberto Kolter. Isolation, characterization, and aggregation of a structured bacterial matrix precursor. Journal of Biological Chemistry, 288:17559–17568, 6 2013. ISSN 00219258. doi: 10.1074/jbc.M113.453605.

15. Anne Diehl, Yvette Roske, Linda Ball, Anup Chowdhury, Matthias Hiller, Noel Molière, Regina Kramer, Daniel Stöppler, Catherine L. Worth, Brigitte Schlegel, Martina Leidert, Nils Cremer, Natalja Erdmann, Daniel Lopez, Heike Stephanowitz, Eberhard Krause, Barth-Jan van Rossum, Peter Schmieder, Udo Heinemann, Kürsşad Turgay, Ümit Akbey, and Hartmut Oschkinat. Structural changes of tasa in biofilm formation of bacillus subtilis. Proceedings of the National Academy of Sciences, 115:3237–3242, 3 2018. ISSN 0027-8424. doi: 10.1073/pnas.1718102115.

16. Mnar Ghrayeb, Shahar Hayet, Neta Lester-Zer, Yael Levi-Kalisman, and Liraz Chai. Fibrilar polymorphism of the bacterial extracellular matrix protein tasa. Microorganisms, 9:529, 3 2021. ISSN 2076-2607. doi: 10.3390/microorganisms9030529.

17. D.N. Azulay, M. Ghrayeb, I.B. Ktorza, I. Nir, R. Nasser, Y.S. Harel, and L. Chai. Colloidal-like aggregation of a functional amyloid protein. Physical Chemistry Chemical Physics, 22, 2020. ISSN 14639076. doi: 10.1039/d0cp03265d.

18. David N. Azulay, Oliver Spaeker, Mnar Ghrayeb, Michaela Wilsch-Bräuninger, Ernesto Scoppola, Manfred Burghammer, Ivo Zizak, Luca Bertinetti, Yael Politi, and Liraz Chai. Multiscale x-ray study of bacillus subtilis biofilms reveals interlinked structural hierarchy and elemental heterogeneity. Proceedings of the National Academy of Sciences, 119, 1 2022. ISSN 0027-8424. doi: 10.1073/pnas.2118107119.

19. Mike Sleutel, Ravi R. Sonani, Jessalyn G. Miller, Fengbin Wang, Andres Gonzalez Socorro, Yang Chen, Reece Martin, Borries Demeler, Michael J. Rudolph, Vikram Alva, Han Remaut, Edward H. Egelman, and Vincent P. Conticello. Donor strand complementation and calcium ion coordination drive the chaperone-free polymerization of archaeal cannulae. Nature Communications, 16:9082, 10 2025. ISSN 2041-1723. doi: 10.1038/s41467-025-64120-8.

20. Mike Sleutel, Adrià Sogues, Andres Gonzalez Socorro, Marcus Fislage, Vikram Alva, Han Remaut, and Vincent P. Conticello. From fibril to framework: P. abyssi abpx illuminates a calcium-responsive family of microbial biomatrix proteins that form thermostable hydrogels. 1 2026. doi: 10.64898/2026.01.04.697558.

21. Nir Ido, Amir Lybman, Shahar Hayet, David N. Azulay, Mnar Ghrayeb, Sajeda Liddawieh, and Liraz Chai. Bacillus subtilis biofilms characterized as hydrogels. insights on water uptake and water binding in biofilms. Soft Matter, 16:6180–6190, 2020. ISSN 1744-683X. doi: 10.1039/D0SM00581A.

22. Silvia Cometta, Dietmar W. Hutmacher, and Liraz Chai. In vitro models for studying implant-associated biofilms - a review from the perspective of bioengineering 3d microenvironments. Biomaterials, 309:122578, 9 2024. ISSN 01429612. doi: 10.1016/j.biomaterials.2024.122578.

23. Matt D.G. Hughes, Daniel West, Rebecca Wurr, Sophie Cussons, Kalila R. Cook, Najet Mahmoudi, David Head, David J. Brockwell, and Lorna Dougan. Competition between cross-linking and force-induced local conformational changes determines the structure and mechanics of labile protein networks. Journal of Colloid and Interface Science, 678:1259–1269, 1 2025. ISSN 00219797. doi: 10.1016/j.jcis.2024.09.183.

24. Sukulya Bunuasunthon, Masahiko Nakamoto, Voravee P. Hoven, and Michiya Matsusaki. Construction of tough hydrogel cross-linked via ionic interaction by protection effect of hydrophobic domains. ACS Biomaterials Science and Engineering, 10:4245–4258, 7 2024. ISSN 2373-9878. doi: 10.1021/acsbiomaterials.4c00681.

25. Jingwen Zhao, Koichi Mayumi, Costantino Creton, and Tetsuharu Narita. Rheological properties of tough hydrogels based on an associating polymer with permanent and transient crosslinks: Effects of crosslinking density. Journal of Rheology, 61:1371–1383, 11 2017. ISSN 0148-6055. doi: 10.1122/1.4997589.

26. Pragnesh N. Dave, Pradip M. Macwan, Bhagvan Kamaliya, and Arvind Kumar. Rheological investigations and swelling behavior of gum ghatti-cl-poly(acrylic acid) hydrogel reinforced with graphene oxide. Journal of Materials Science: Materials in Engineering, 20:11, 2 2025. ISSN 3004-8958. doi: 10.1186/s40712-024-00148-w.

27. Amra Noa, Hui-Shun Kuan, Vera Aschmann, Vasily Zaburdaev, and Lennart Hilbert. The hierarchical packing of euchromatin domains can be described as multiplicative cascades. PLOS Computational Biology, 17:e1008974, 5 2021. ISSN 1553-7358. doi: 10.1371/journal.pcbi.1008974.

28. Hadrien Salat, Roberto Murcio, and Elsa Arcaute. Multifractal methodology. Physica A: Statistical Mechanics and its Applications, 473:467–487, 5 2017. ISSN 03784371. doi: 10.1016/j.physa.2017.01.041.

29. J. Teixeira. Small-angle scattering by fractal systems. Journal of Applied Crystallography, 21:781–785, 12 1988. ISSN 0021-8898. doi: 10.1107/S0021889888000263.

30. Byong kyu Shin and Sunil Saxena. Insight into potential cu(ii)-binding motifs in the four pseudorepeats of tau protein. The Journal of Physical Chemistry B, 115:15067–15078, 12 2011. ISSN 1520-6106. doi: 10.1021/jp204410h.

31. Byong kyu Shin and Sunil Saxena. Substantial contribution of the two imidazole rings of the his13-his14 dyad to cu(ii) binding in amyloid-beta (1-16) at physiological ph and its significance. The Journal of Physical Chemistry A, 115:9590–9602, 9 2011. ISSN 1089-5639. doi: 10.1021/jp200379m.

32. Stefan Stoll and Arthur Schweiger. Easyspin, a comprehensive software package for spectral simulation and analysis in epr. Journal of Magnetic Resonance, 178:42–55, 1 2006. ISSN 10907807. doi: 10.1016/j.jmr.2005.08.013.

33. J. Peisach and W.E. Blumberg. Structural implications derived from the analysis of electron paramagnetic resonance spectra of natural and artificial copper proteins. Archives of Biochemistry and Biophysics, 165:691–708, 12 1974. ISSN 00039861. doi: 10.1016/0003-9861(74)90298-7.

34. A Artemenko, A Shchukarev, P štenclová, T Wågberg, J Segervald, X Jia, and A Kromka. Reference xps spectra of amino acids. IOP Conference Series: Materials Science and Engineering, 1050:012001, 1 2021. ISSN 1757-8981. doi: 10.1088/1757-899X/1050/1/012001.

35. Paul S Bagus, Carmen Sousa, and Francesc Illas. Xps binding energy shifts as a function of bond distances: a case study of co. Journal of Physics: Condensed Matter, 34:154004, 4 2022. ISSN 0953-8984. doi: 10.1088/1361-648X/ac4dc0.

36. J.F. Moulder and J. Chastain. Handbook of X-ray Photoelectron Spectroscopy: A Reference Book of Standard Spectra for Identification and Interpretation of XPS Data. Physical Electronics Division, Perkin-Elmer Corporation, 1992. ISBN 9780962702624.

37. Milan Remko, Daniel Fitz, and Bernd Michael Rode. Effect of metal ions (li+, na+, k+, mg2+, ca2+, ni2+, cu2+ and zn2+) and water coordination on the structure and properties of l-histidine and zwitterionic l-histidine. Amino Acids, 39:1309–1319, 11 2010. ISSN 0939-4451. doi: 10.1007/s00726-010-0573-8.

38. Tianxin Fu, Shu Zhang, Yanan Sheng, Yuchao Feng, Yingjun Jiang, Yiwei Zhang, Miao Yu, and Changyuan Wang. Isolation and characterization of zinc-binding peptides from mung bean protein hydrolysates. European Food Research and Technology, 246:113–124, 1 2020. ISSN 1438-2377. doi: 10.1007/s00217-019-03397-8.

39. Jiajing Zhang, Wei Yang, Jean-Philip Piquemal, and Pengyu Ren. Modeling structural coordination and ligand binding in zinc proteins with a polarizable potential. Journal of Chemical Theory and Computation, 8:1314–1324, 4 2012. ISSN 1549-9618. doi: 10.1021/ct200812y.

40. Craig K. Esser, Robert L. Bugianesi, Charles G. Caldwell, Kevin T. Chapman, Philippe L. Durette, Narindar N. Girotra, Ihor E. Kopka, Thomas J. Lanza, Dorothy A. Levorse, Malcolm MacCoss, Karen A. Owens, Mitree M. Ponpipom, Joseph P. Simeone, Richard K. Harrison, Lisa Niedzwiecki, Joseph W. Becker, Alice I. Marcy, Melinda G. Axel, Amy J. Christen, Joseph McDonnell, Vernon L. Moore, Julie M. Olszewski, Cheryl Saphos, Denise M. Visco, Frank Shen, Adria Colletti, Philip A. Krieter, and William K. Hagmann. Inhibition of stromelysin-1 (mmp-3) by p <sub>1</sub> ‘-biphenylylethyl carboxyalkyl dipeptides. Journal of Medicinal Chemistry, 40:1026–1040, 3 1997. ISSN 0022-2623. doi: 10.1021/jm960465t.

41. Theyencheri Narayanan, Michael Sztucki, Thomas Zinn, Jérôme Kieffer, Alejandro Homs-Puron, Jacques Gorini, Pierre Van Vaerenbergh, and Peter Boesecke. Performance of the time-resolved ultra-small-angle x-ray scattering beamline with the extremely brilliant source. Applied Crystallography, 55(1):98–111, 2022.

42. Theyencheri Narayanan, Michael Sztucki, Pierre Van Vaerenbergh, Joachim Léonardon, Jacques Gorini, Laurent Claustre, Franc Sever, John Morse, and Peter Boesecke. A multipurpose instrument for time-resolved ultra-small-angle and coherent X-ray scattering. Journal of Applied Crystallography, 51(6):1511–1524, Dec 2018. doi: 10.1107/S1600576718012748.

43. Avi Ginsburg, Tal Ben-Nun, Roi Asor, Asaf Shemesh, Israel Ringel, and Uri Raviv. Reciprocal grids: A hierarchical algorithm for computing solution x-ray scattering curves from supramolecular complexes at high resolution. Journal of Chemical Information and Modeling, 56:1518–1527, 8 2016. ISSN 1549-9596. doi: 10.1021/acs.jcim.6b00159.

44. Avi Ginsburg, Asaf Shemesh, Abigail Millgram, Raviv Dharan, Yael Levi-Kalisman, Israel Ringel, and Uri Raviv. Structure of dynamic, taxol-stabilized, and gmppcp-stabilized microtubule. The Journal of Physical Chemistry B, 121:8427–8436, 9 2017. ISSN 1520-6106. doi: 10.1021/acs.jpcb.7b01057.

45. Avi Ginsburg, Tal Ben-Nun, Roi Asor, Asaf Shemesh, Lea Fink, Roee Tekoah, Yehonatan Levartovsky, Daniel Khaykelson, Raviv Dharan, Amos Fellig, and Uri Raviv. D+: software for high-resolution hierarchical modeling of solution x-ray scattering from complex structures. Journal of Applied Crystallography, 52(1):219–242, 2019.

46. Moshe Nadler, Ariel Steiner, Tom Dvir, Or Szekely, Pablo Szekely, Avi Ginsburg, Roi Asor, Roy Resh, Carmen Tamburu, Menahem Peres, et al. Following the structural changes during zinc-induced crystallization of charged membranes using time-resolved solution xray scattering. Soft Matter, 7(4):1512–1523, 2011.

