## Supplementary Information for "A bacterial extracellular matrix protein forms a supramolecular metallogel"

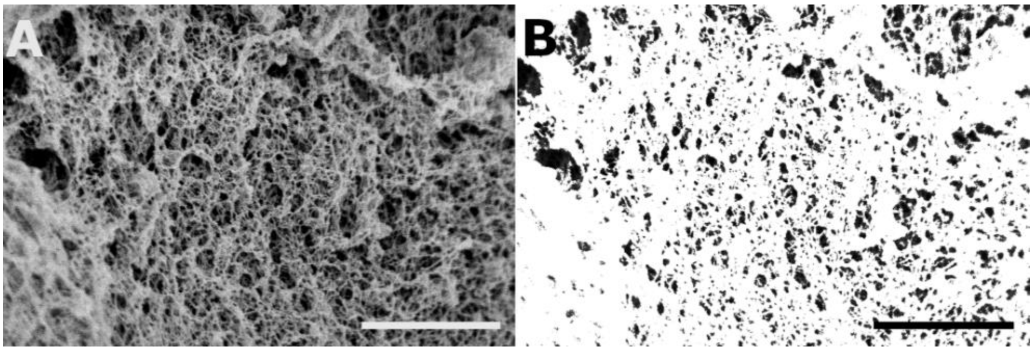


**Fig. S1. SEM image-based analysis for pore area.** (A) An example of an SEM image of 12 µM TasA, 20 mM $ZnCl_2$ hydrogel sample used for pore area analysis. (B) The same image in A after threshold adjustment using ImageJ; pores are shown in black and gel network in white. Scale bar is 1 µm.

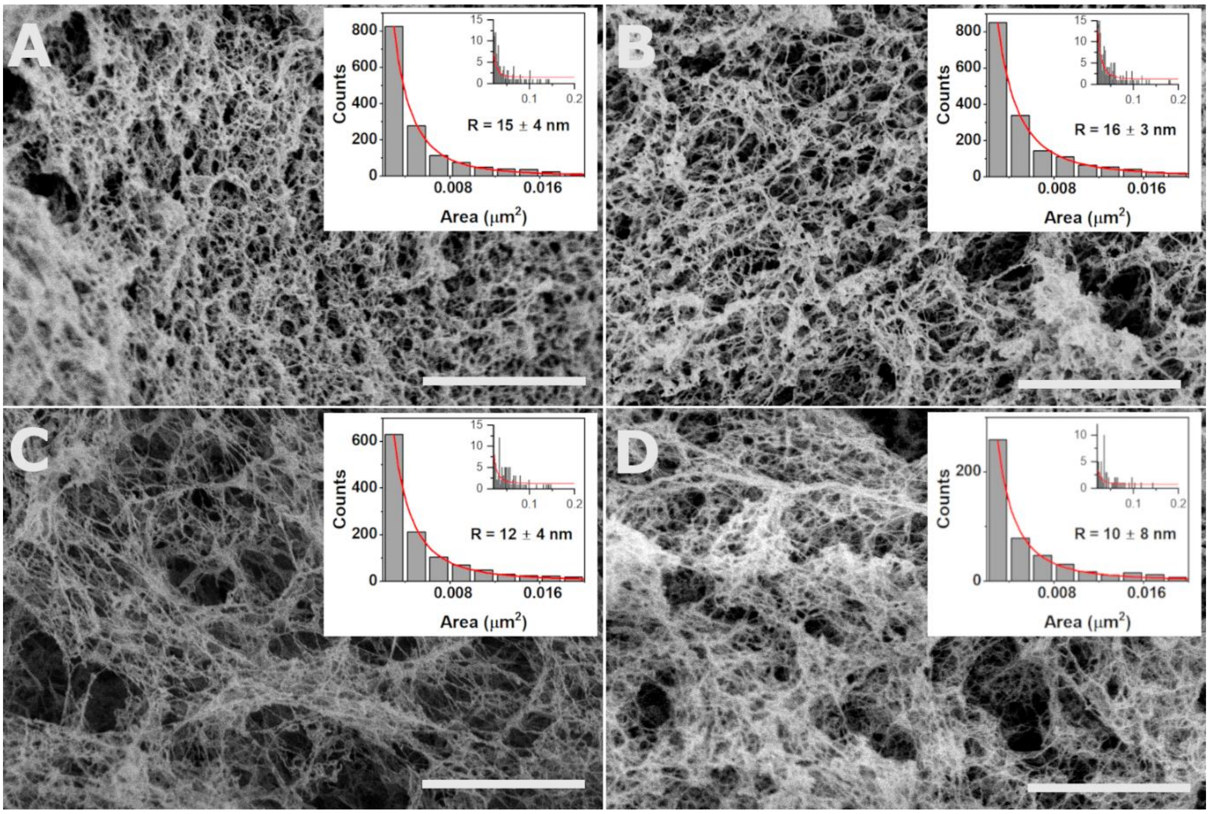


**Fig. S2. SEM images of TasA-Zn hydrogels and pore area analysis.** (A) TasA-Zn hydrogel made by mixing: 12 µM TasA with 20 mM $ZnCl_2$, (B) 20 µM TasA with 20 mM $ZnCl_2$, (C) 12 µM with TasA 100 mM $ZnCl_2$, and (D) 20 µM TasA with 100 mM $ZnCl_2$. Inset for each gelation condition shows the area distribution of the pores, with zoom into the lower values denoted by the distributions at the top right of the inset. Data are averaged over N=11 images (A), N=12 images (B), N=8 images (C), and N=5 images (D). Red lines show a Gaussian fit which gives the mean pore area *A* at each gelation condition, from which the radius was calculated assuming a circular pore geometry, using $A = \pi R^2$. Calculated radii are indicated on respective plots. Scale bars in SEM images depict 1 µm.

**Table S1:** Pore radii and area as analyzed using SEM images, such as those shown in Fig. S2.

| | R (nm) | Pore Area ($nm^2$) |
|---|---|---|
| **12 μM TasA 20 mM $ZnCl_2$ (A)** | $15 \pm 4$ | $717 \pm 233$ |
| **20 μM TasA 20 mM $ZnCl_2$ (B)** | $16 \pm 3$ | $755 \pm 201$ |
| **12 μM TasA 100 mM $ZnCl_2$ (C)** | $12 \pm 4$ | $437 \pm 228$ |
| **20 μM TasA 100 mM $ZnCl_2$ (D)** | $10 \pm 8$ | $304 \pm 348$ |

**Table S2:** Optical density values of TasA-Zn solutions after 18 hours of incubation at room temperature. Samples with optical density (OD) values larger than 0.12 considered as gels.

| [TasA] (μM) | [$ZnCl_2$] (mM) | $OD_{600nm}$@18h (A.U.) |
|---|---|---|
| 4 | 0 | 0.099 ±0.002 |
| 4 | 0.5 | 0.104±0.003 |
| 4 | 1 | 0.114±0.001 |
| 4 | 3 | 0.125±0.002 |
| 4 | 5 | 0.139±0.006 |
| 4 | 10 | 0.14±0.01 |
| 4 | 20 | 0.152±0.004 |
| 4 | 50 | 0.148±0.007 |
| 4 | 100 | 0.153±0.001 |
| 12 | 0 | 0.109±0.002 |
| 12 | 0.5 | 0.101±0.001 |
| 12 | 1 | 0.13±0.03 |
| 12 | 3 | 0.14±0.02 |
| 12 | 5 | 0.116±0.002 |
| 12 | 10 | 0.162±0.004 |
| 12 | 20 | 0.158±0.007 |
| 12 | 50 | 0.135±0.001 |
| 12 | 100 | 0.12±0.01 |
| 20 | 0 | 0.114±0.005 |
| 20 | 0.5 | 0.095±0.001 |
| 20 | 1 | 0.096±0.001 |
| 20 | 3 | 0.16±0.05 |
| 20 | 5 | 0.14±0.01 |
| 20 | 10 | 0.169±0.008 |
| 20 | 20 | 0.123±0.005 |
| 20 | 50 | 0.17±0.03 |
| 20 | 100 | 0.18±0.03 |

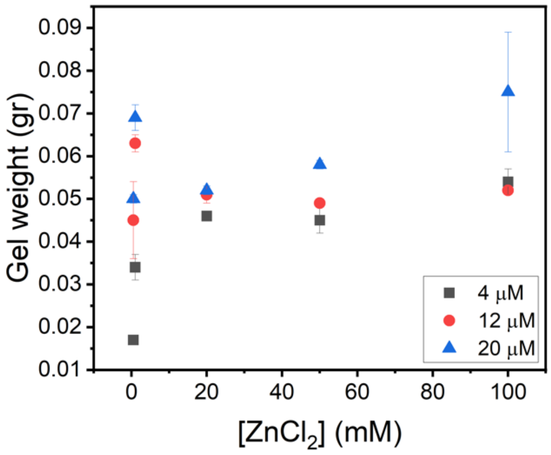


**Fig. S3. TasA-Zn hydrogel weight as function of $ZnCl_2$ concentration added to TasA at 4 µm, 12µm, and 20 µm.** The weight of TasA-Zn gel, formed by mixing TasA in concentrations as indicated in the figure legend (4 µM (black squares), 12 µM (red circles), and 20 µM (blue triangles)) with increasing concentrations of $ZnCl_2$ solutions. Error bars represent the standard deviation of three replicate measurements. Increasing $ZnCl_2$ concentration leads to higher gel weight, until saturation is reached at ~ 20 mM $ZnCl_2$.

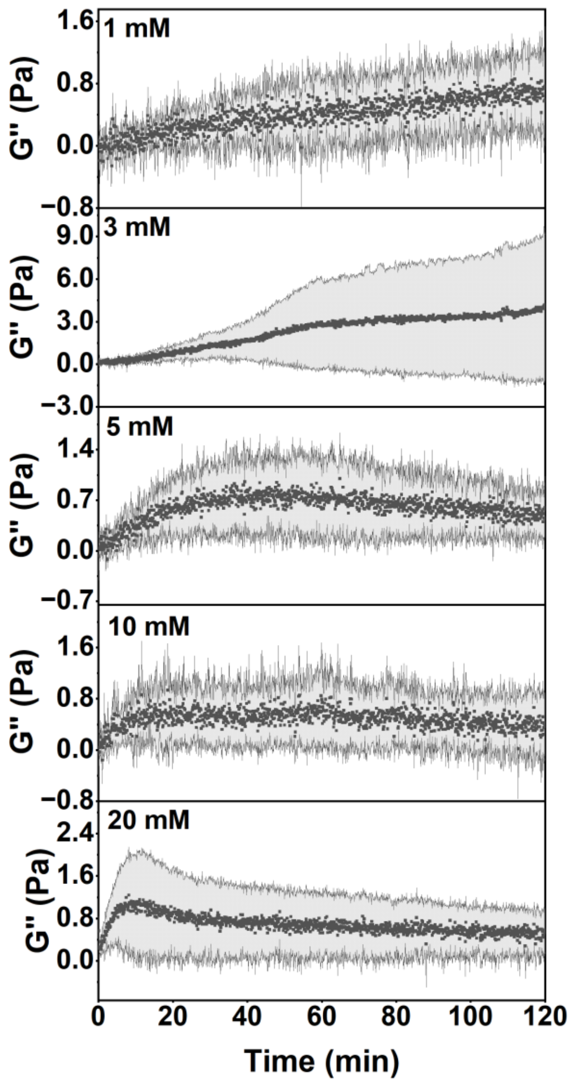


**Fig. S4. Rheology of TasA-Zn aggregates in solution.** Rheology measurements showing the loss modulus (G'') of a TasA-Zn gel formed with 4 µM TasA and increasing concentrations of $ZnCl_2$ in the range 1 mM - 20 mM, plotted as a function of time. Average values are denoted by dark symbols, and the error is denoted by grey area.

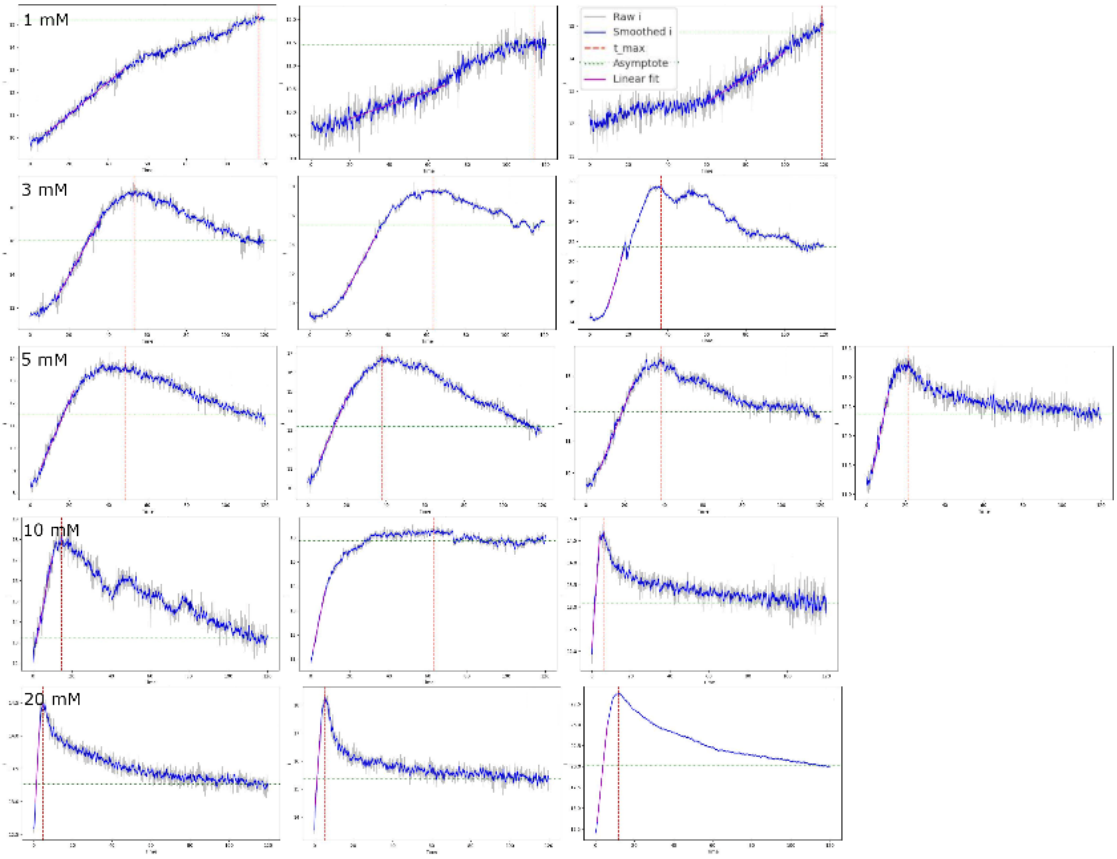


**Fig. S5. Analysis of rheology measurements of TasA-Zn aggregates in solution.** The storage modulus of 4 µM TasA with increasing concentrations of $ZnCl_2$ in the range of 1-20 mM was plotted as a function of time (black). Data was smoothed (blue), and linearly fitted (purple), the slope of linear fit corresponds to K (increase rate) and $G'_{max}$ (maximal G' value) corresponds to $t_{max}$.

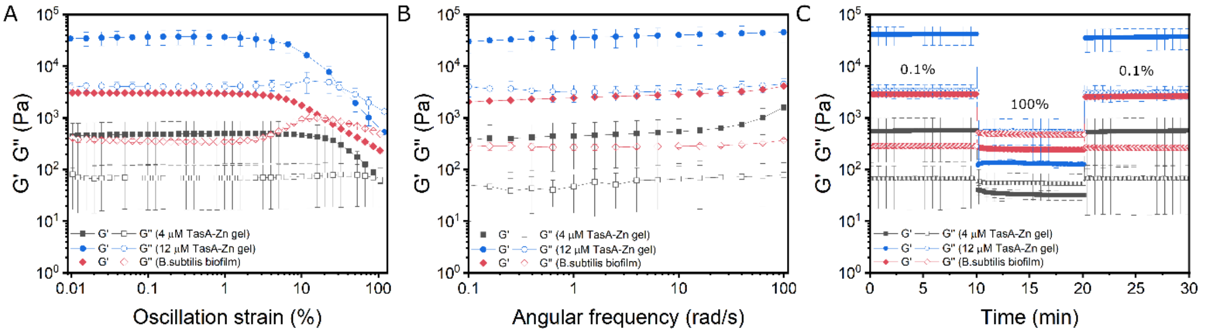


**Fig. S6. Rheological characterization of TasA-Zn hydrogels and a wild-type *B. subtilis* biofilm.** (A) Strain sweep, (B) frequency sweep, and (C) time sweep measurements of G' and G'' of TasA-Zn gels and *B. subtilis* biofilm. Storage modulus (G', full symbols) and loss modulus (G'', empty symbols) are plotted as function of oscillation strain in the range of 0.01-100% for hydrogels made from 4 µM TasA and 20 mM $ZnCl_2$ (black), 12 µM TasA and 20 mM $ZnCl_2$ (blue), and WT *B. subtilis* biofilm (red). Data are averaged over three replicates.

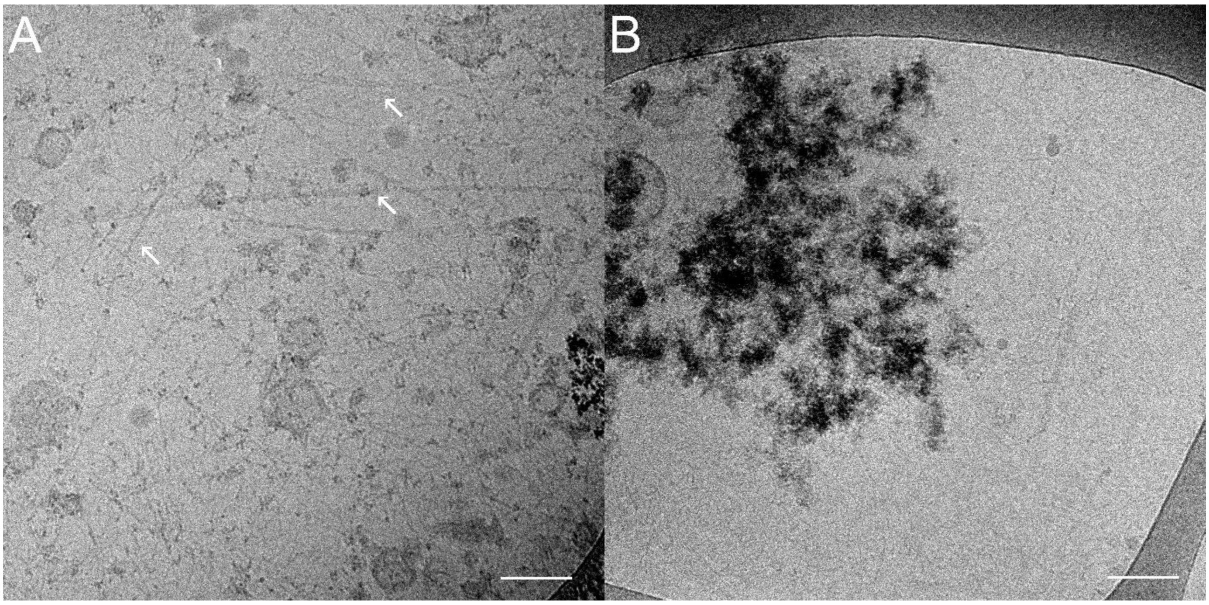


**Fig. S7. TasA forms different morphologies with low $ZnCl_2$ concentration.** Cryo-TEM images of 12 µM TasA with 1 mM $ZnCl_2$, showing the coexistence of fibers and bundles (the latter are indicated with arrows) (A) and clusters (B). Scale bar corresponds to 100 nm.

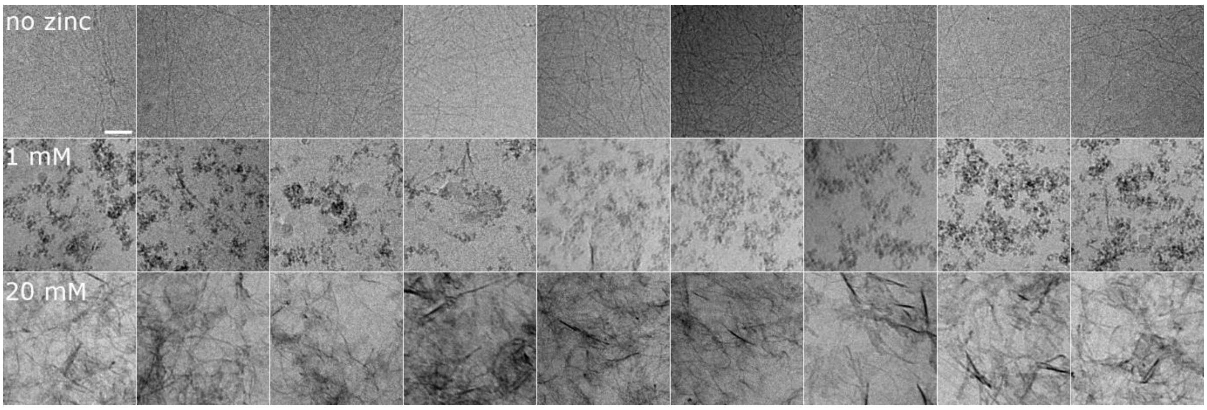


**Fig. S8. TEM images used to analyze the multifractality of TasA in different $ZnCl_2$ concentrations.** TEM regions of interest (ROI) of 12 µM TasA with no zinc (upper row), 1 mM $ZnCl_2$ (middle row), and 20 mM $ZnCl_2$ (bottom row). Scale bar corresponds to 200 nm.

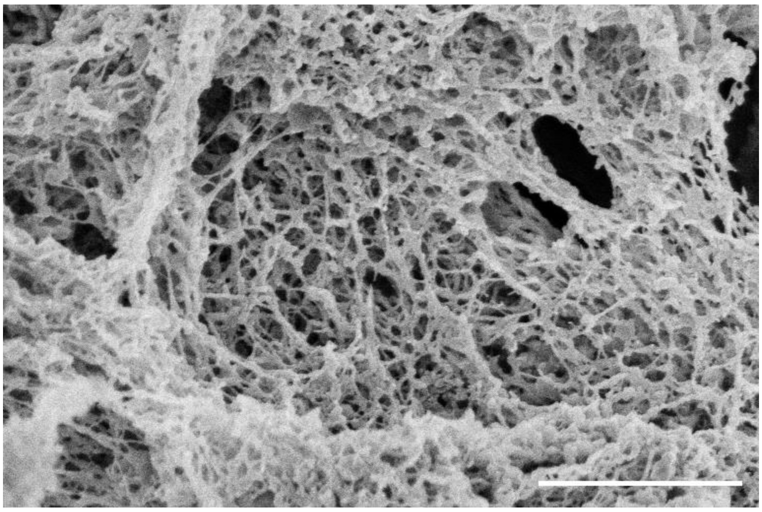

**Fig. S9. TasA forms hydrogel with $CuCl_2$.** Scanning electron microscopy image of 12 µM TasA with 20 mM $CuCl_2$ showing the formation of 3D network. Scale bar corresponds to 1 µm.

**Table S3:** A summary of EPR fitting parameters

| | Tris buffer | TasA | TasA – Zn gel |
|---|---|---|---|
| **g ⊥** | 2.06 | 2.06 | 2.06 |
| **g ǁ** | 2.22 | 2.28 | 2.27 |
| **A ⊥ (MHz)** | 50 | 50 | 50 |
| **A ǁ (MHz)** | 580 | 410 | 470 |

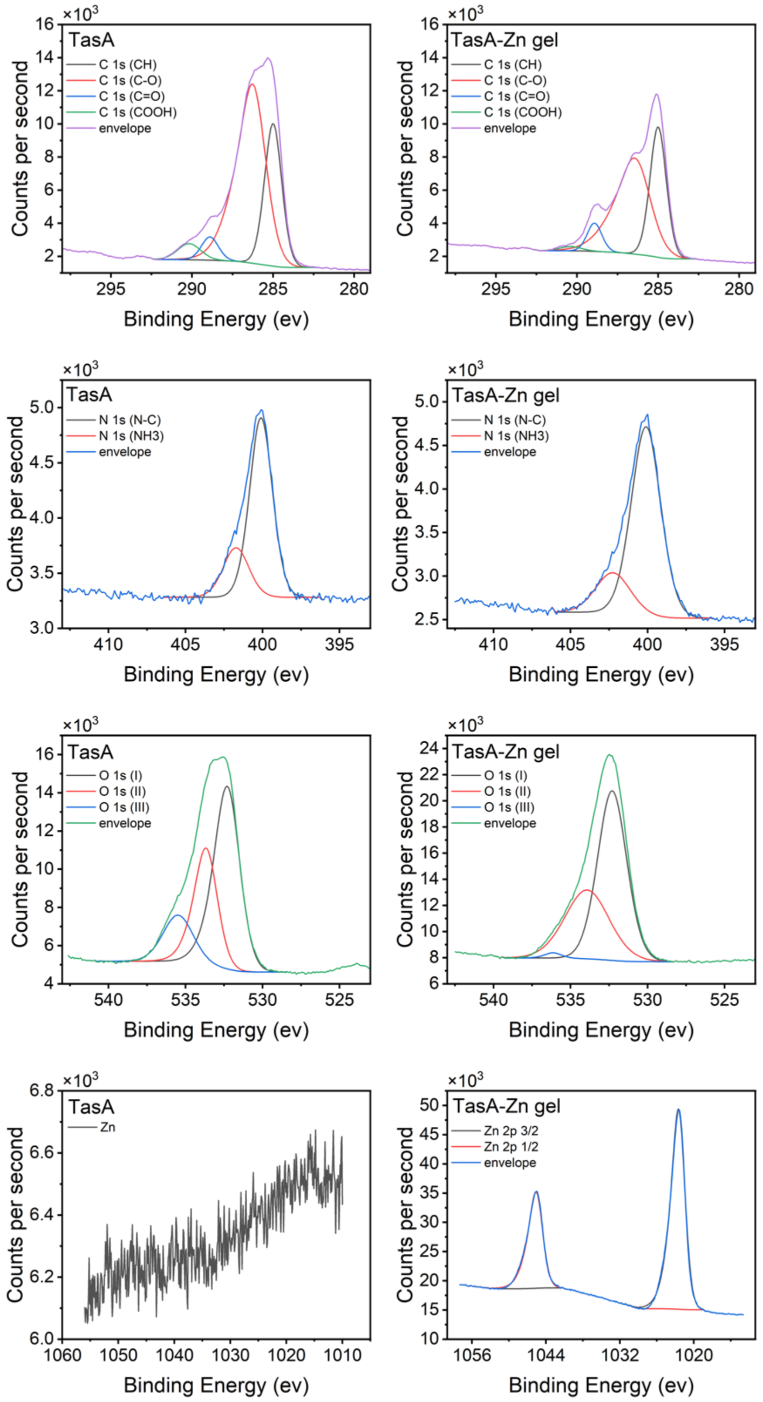


**Fig. S10. X-ray photoelectron spectroscopy (XPS) analysis of TasA and TasA-Zn hydrogel.** XPS spectra of the C 1s, N 1s, O 1s, and Zn 2p regions for TasA (left column) and TasA/Zn gel (right column). The C 1s spectra were deconvoluted into contributions from C–C/C–H, C–O, C=O, and O–C=O functional groups. The N 1s spectra show components attributed to N–C and protonated amine ($NH_3^+$) species. The O 1s region was fitted with three oxygen components corresponding to different oxygen-containing environments. In the TasA-Zn hydrogel sample, additional Zn $2p_{3/2}$ and Zn $2p_{1/2}$ peaks were observed, confirming the presence of $Zn^{2+}$ ions in the gel. The envelopes represent the overall fitted spectra obtained from the sum of individual components.

**Table S4:** Binding energies (eV) from XPS data assigned to functional groups

| | TasA | TasA/Zn |
|---|---|---|
| **CH** | 285 $\pm$ 0 | 285 $\pm$ 0 |
| **C − O** | 286.4 $\pm$ 0.1 | 286.3 $\pm$ 0.1 |
| **C = O** | 288.6 $\pm$ 0.4 | 288.3 $\pm$ 0.1 |
| **COOH** | 289.8 $\pm$ 0.6 | 289.7 $\pm$ 0.3 |
| **N − C** | 400.1 $\pm$ 0.0 | 400.1 $\pm$ 0.0 |
| **$NH_3$** | 401.7 $\pm$ 0.1 | 402.2 $\pm$ 0.1 |
| **O(I)** | 532.1 $\pm$ 0.3 | 532.1 $\pm$ 0.2 |
| **O(II)** | 533.5 $\pm$ 0.3 | 533.4 $\pm$ 0.5 |
| **O(III)** | 535.6 $\pm$ 0.1 | 536.1 $\pm$ 0.1 |
| **Zn 2p 3/2** | ___ | 1022.6 $\pm$ 0.1 |
| **Zn 2p 1/2** | ___ | 1045.7 $\pm$ 0.1 |

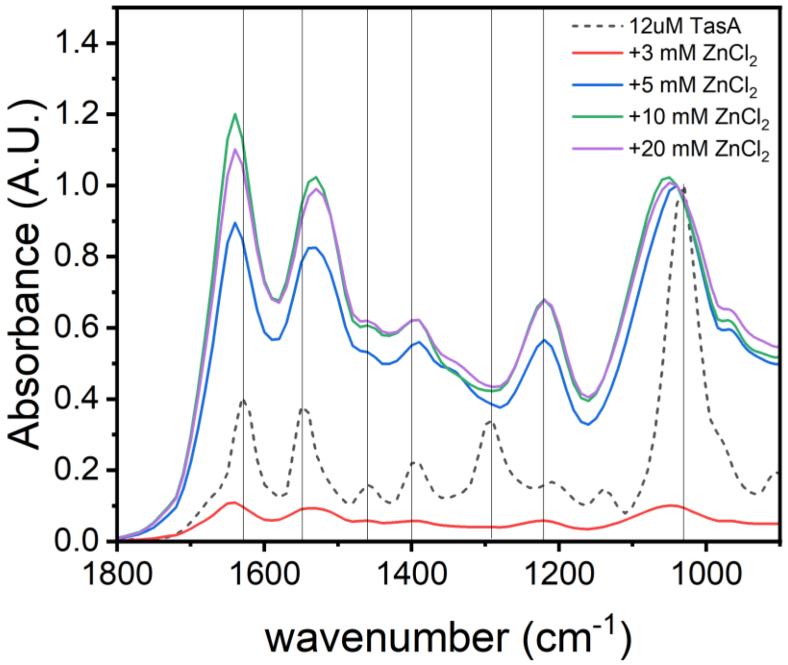


**Fig. S11. Effect of zinc ions addition to TasA as observed by Fourier transform Infrared (FTIR) spectroscopy.** FTIR spectrum of 12 µM TasA (no added zinc chloride) (black), and with the addition of zinc chloride solutions, according to the concentrations specified in the legend (colours), showing a clear shift in peak positions of Amide I, II, and III regions.

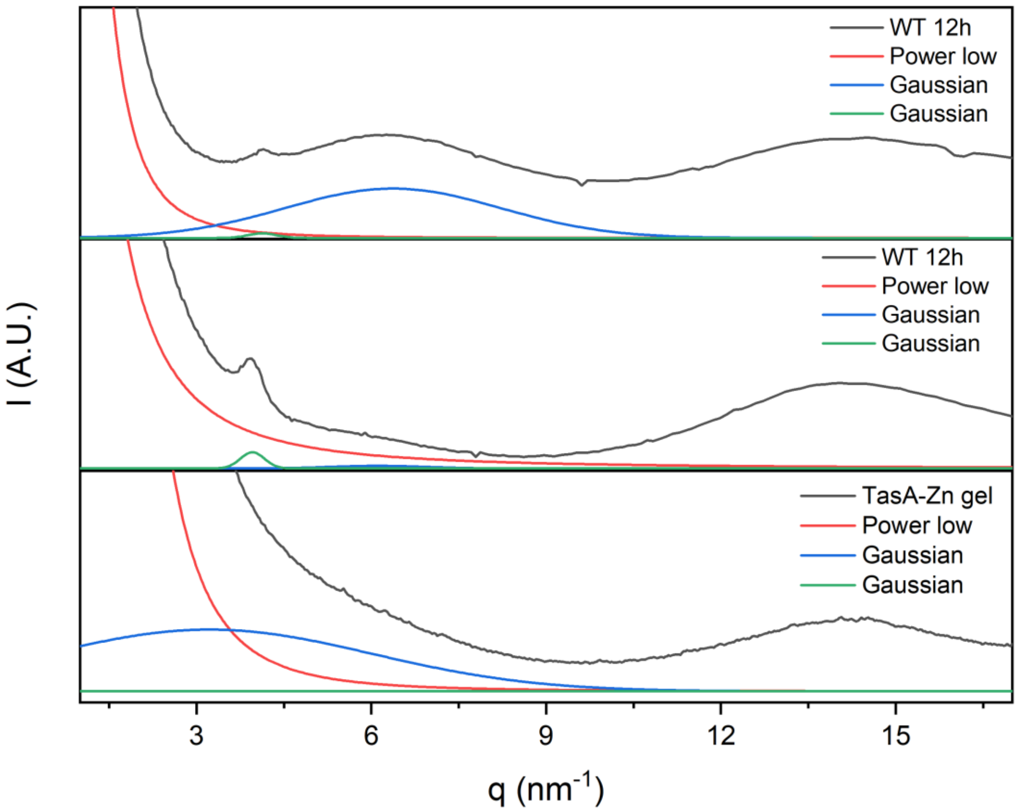


**Fig. S12. X-ray diffraction (XRD) spectra and fitting of TasA–Zn gel and WT biofilm samples.** Experimental XRD profiles of TasA–Zn gel (bottom panel) (black) is shown together with the corresponding fits (color) that we used to deconvolve the ~ 6 $nm^{-1}$ peak, as follows: the scattering curves were fitted using a combination of a low-q power-law contribution (red), and Gaussian peaks (blue and green), representing characteristic structural correlations. In WT biofilm samples ('WT12h', black curves in the middle and top panels), a broad peak centered at ~ 6 $nm^{-1}$ is observed in some cases (top) and absent in others (middle). WT 12 h signals are taken from Azulay et al., PNAS (2022). This feature is absent in the TasA–Zn gel spectrum.

### Experimental

<u>Native TasA purification</u>

TasA was purified natively as described previously[1,2]. Briefly, a preculture of *B.subtilis* double mutant ΔsinR-ΔEPS was diluted 1:100 in MSgg Broth[3]. Bacterial culture centrifuged (10,000xg, 15 min, 4°C) and pellets were sonicated in 1.5 M saline extraction buffer (1.5 M NaCl, 2 mM $MgCl_2$, 5 mM Potassium phosphate pH 7.0, 0.1 M MOPS pH 7.0), then stirred for 30 min at room temperature. After another centrifugation (10,000xg, 15 min, 4°C) supernatant was filtered through 0.45 µm filters and kept at 4°C for 3 days. The crude extract was then centrifuged (17,000xg, 15 min, 4°C) and supernatant was loaded on HiLoad 26/60 Superdex S200 sizing column that was pre-equilibrated with 20 mM Tris pH 8.0. Fractions of pure TasA were eluted and kept at -20°C until used. TasA was concentrated using Amicon centrifugal filter tubes with MWCO 10 KDa. Final concentration of TasA samples were determined using BCA assay.

<u>TasA-Zn gel samples</u>

TasA was mixed with $ZnCl_2$ (using a X100 stock solution) then sample was centrifuged 4600xg, 5 min to form a gel. Gel was collected for further analysis.

TasA-Zn gel weight

4, 12, and 20 µM TasA was mixed with 100X $ZnCl_2$ solution to reach final concentrations of 1-100 mM. Solutions were centrifuged 4600xg, 5 min. Gels weighed with analytical weight.

TasA-Zn gel formation and Bottom-up tube experiment

A 1 mL of 12 µM TasA was mixed with 100X $ZnCl_2$ solution to reach final concentrations of 1 and 20 mM $ZnCl_2$ solutions and then centrifuged 4600xg, 5 min, 4°C. Images of the tube were taken with Nikon Camera.

Scanning Electron microscopy

1 mL of TasA (12, and 20 µM) was mixed with 100X $ZnCl_2$ solution to reach final concentrations of 20, and 100 mM and gels were prepared as described above. Gels were collected and washed with triple distilled water for 16 hours, then washed with gradient concentration of ethanol solution for 10 min for each solution and kept in 100% ethanol for another 16 hours. Later samples were dried using the supercritical drying with $CO_2$ using the Tousimis manual critical point dryer, SAMDRI-PVT-3D. The drying prosses consists of pressuring the chamber and rinsing the gels with liquid $CO_2$ once for 7 min. Then heating to 38 °C and slowly depressurization. Samples were coated with Iridium for 60 sec using the Quorum coating machine and scanned with extra-resolution scanning electron microscope (XHR Magellan 400 L). Similar preparation was done for 12 µM TasA with 20 mM $CuCl_2$ hydrogel.

Pore size analysis

SEM images were analyzed using image J as follows: images were converted to 8 bit, and a threshold was applied to differentiate between pores and gel. We used images with 40,000× or 80,000× magnifications, in order to sample a significant number of pores per image. Thresholding images with these resolutions introduced outlier pores of the order of a few pixels that appeared to be smaller than their actual size. This was corrected by setting a lower bound of 10 pixels to all images so that pores, smaller than 10 pixels, were excluded from our analysis. This set the lower limit in our analysis to pore areas larger than 2000 $nm^2$.

Cryogenic Transmission Electron Microscopy

TasA-Zn samples were prepared as explained above. Gel samples were washed for 16 hours. A 3 µL droplet of TasA solution or gel was deposited on a glow-discharged TEM grid (300 mesh Cu Lacey substrate grid; Ted Pella). The excess liquid was blotted off with filter paper, and the specimen was rapidly plunged into liquid ethane pre-cooled with liquid nitrogen in a controlled environment using a Vitrobot Mark IV (FEI). The vitrified samples were transferred to a cryo-specimen holder (Gatan 626) and examined at −177 °C using a FEI Tecnai 12 $G^2$ TWIN TEM instrument operated at 120 kV in low-dose mode. Grids were imaged at a few micrometers under focus to increase phase contrast. The images were recorded with a 4K × 4K FEI Eagle charged-couple device (CCD) camera. Similar preparation was done for samples measure by small angle X-ray scattering (SAXS).

Fractal dimension analysis

*Image analysis*

TEM images of aggregates were classified according to 3 different TasA and ZnCl concentrations, yielding a total of 9 unique sets. For each concentration set, images with the same magnification were selected and a total of 9 1000 pixels × 1000 pixels regions of interest (ROIs) were selected by visual inspection to be mostly covered by aggregates (see Fig. S8 as an example).

Primary image analysis was performed using Matlab. Given that aggregates in the original images show up as dark objects against a light background, the ROIs were inverted as the first step of the pipeline, to ensure that structures of interest in the images are bright. Multi-fractal analysis and structure factor calculations were then performed on each of the 9 *per-condition* ROIs. Plots of all quantifiers for each condition are presented as the mean over the 9 ROIs for that condition, with the error bars displayed as ±1 SEM.

*Multifractal analysis*

We employ the standard box-counting method for calculating the multifractal spectra of the aggregates[4] as we have previously done for the quantification of the DNA structure in the cell nucleus[5]. For box-counting, a square grid of grid size $r \times r$ pixels is laid over the image to be analysed. As multifractal analysis is performed on measures, the intensity $I_i$ in the $i^{th}$ box is converted into a probability measure $P_i$ via normalization by the total image intensity $I_{tot}$: $P_i = I_i/I_{tot}$. For the actual calculation of the spectra, we need the generalized moments of the probability distribution $\mu^q{}_i$, which can be obtained by raising $P_i$ to the exponents $q$:

$$\mu_i^q = \frac{(P_i)^q}{\sum_i (P_i)^q} \tag{1}$$

From the intensity probability distribution and it's generalized moments, we can calculate the generalized dimension $D_q$ for different $q$'s. We also quantify the so-called singularity strength $\alpha$ or the scaling exponent exhibited by a subset of an image and the respective fractal dimension of this subset $f(\alpha)$[4,6] to produce the multifractal spectrum of the system. Together these quantities characterize the multifractal character of the TasA aggregates. For this, we first calculate the quantity $\tau(q)$, defined as the following limit:

$$\tau(q) = \lim_{r \to 0} \frac{\log\left(\sum_i \mu_i^q(r)\right)}{\log r} \tag{2}$$

Using $\tau(q)$, the rest of the quantities are defined as follows:

$$D_q = \frac{\tau(q)}{(q-1)} \tag{3}$$

$$\alpha(q) = \frac{d\tau(q)}{dq} \tag{4}$$

$$f\big(\alpha(q)\big) = \alpha(q)q - \tau(q) \tag{5}$$

To obtain $\tau(q)$ in practice, we calculate $\sum_\mathrm{i} \mu_i^q(r)$ (via Eq. 1) for a set of different grid sizes $r$ and plot $\sum_\mathrm{i} \mu_i^q(r)$ *vs. r* in log-log scale. $\tau(q)$ is then given by the slope of the linear fit to this plot.

*Structure factor*

The structure factor of the aggregates is calculated from the two-dimensional spatial Fourier transform of the selected ROIs. The structure factor is defined as follows:

$$F(k) = \frac{\langle I(k)I(-k)\rangle_k}{I_{tot}^2} \tag{6}$$

where $I(\mathbf{k})$ is the 2D Fourier transform of the image intensities, $\langle ... \rangle_k$ denotes the radial average in the spatial frequency domain, and $I_{tot}$ is the total intensity of the image. $k$ is the spatial frequency in units of $nm^{-1}$, and the conversion from pixel units for the images to $nm^{-1}$ is carried out by measuring the length of scale bars on the original TEM images in terms of pixels.

The structure factor of the aggregates as calculated via this method contains contributions not just from the aggregates themselves, but also possibly from the background. To understand the contribution of the background signal to the structure factor, we carried out structure factor analysis on the background as well for a selected condition. For this, we selected ROIs unobscured by the TEM grid or aggregates. We then calculated the structure factor of these background ROIs as described above. However, we normalized the Fourier transforms of the background intensities by the average over the squares of the total intensities of the aggregate ROIs belonging to the same condition as the background. The formula for the structure factor of the background then becomes:

$$F_{bg}(k) = \frac{\langle I_{bg}(k) I_{bg}(-k) \rangle_k}{\langle I_{tot}^2 \rangle} \quad (7)$$

where $\langle I_{tot}^2 \rangle$ is obtained over the aggregate ROIs, and not the background ROIs themselves.

Rheology measurements of TasA-Zn hydrogel

Gel was prepared using 25 mL of 12 µM TasA solution that was mixed with 25 µL of 2 M $ZnCl_2$ solution (final concentration is 20 mM), and centrifuged 4600xg, 15 min, 4°C. Gel was loaded in the stage of TA rheometer equipped with 20 mm rheometer plate. Temperature was set at 4°C during the measurements. Strain sweep experiment was applied in a shear strain of 0.01-100% with 10 rad/s angular frequency. Sample soaked for 10 sec before measuring. Followed with frequency sweep experiment by applying 0.1% shear strain in an angular frequency range of 0.1-100 rad/sec. Also here, sample was soaked for 10 sec prior to measuring. Finaly, a time sweep experiment was performed to learn about gel self-healing as follows: storage and loss moduli were measured by applying 0.1% strain in frequency of 10 rad/s for 10 min, then strain was increased to 100% for another 10 min and lastly set back to 0.1% for 10 min. Data shown is an average over three repeats.

Rheology of TasA-Zn solutions

500 µL of TasA (4 µM) was mixed with 100X stock solution of $ZnCl_2$ to reach final concentration of 1,3,5,10, and 20 mM. solution was then loaded on the rheometer stage equipped with 20 mm rheometer plate, with loading gap of 1000 µm. Time sweep measurements were performed at constant strain of 0.1% and frequency of 10 rad/sec, at room temperature (25°C). Three replicates were measured for each $ZnCl_2$ concentration.

Rheology of *Bacillus subtilis* biofilms

A 2 µL drop of *Bacillus subtilis* (NCBI 3610) preculture in LB broth (37°C, 250 rpm, 16 h) was placed on an MSgg-agar plate, incubated for 72 h at 30°C. 10 biofilms were collected and mixed to get homogenous sample. Biofilms were measured similarly to TasA-Zn hydrogel samples. Data shown is an average over three repeats.

Thermogravimetric analysis (TGA)

2 mL TasA-Zn hydrogel samples were prepared as described above; gels were washed 16 h with water before the measurement. TasA-Zn hydrogel samples were placed into TGA crucibles. The mass loss was

measured while heating the sample in a temperature range of 25-200°C with 2°C/min heating rate. Data shown is an average of three repeats.

Data analysis and calculations

The weight of the wet gel samples was $W_{wet\ gel} = 41 \pm 2\ mg$, and for the dry gel was $W_{dry\ gel} = 1.2 \pm 0.7\ mg$, thus we calculated the total weight of water withing the sample using the following equation:

$$W_{tot\ water} = W_{wet\ gel} - W_{dry\ gel}$$

$$W_{tot\ water} = 40 \pm 2\ mg$$

the percentage of water in gel sample was then calculated

$$\%total\ water = \frac{W_{tot\ water}}{W_{wet\ gel}} * 100\% = 97\%$$

$$\%e_{tot\ water} = \left(\left(\frac{2}{41}\right)^2 + \left(\frac{2}{40}\right)^2\right)^{0.5} * 97\% = 7\%$$

$$\%total\ water = 97\ \pm 7\ \%$$

Electron paramagnetic resonance (EPR)

$Cu^{2+}$ low-temperature EPR measurements were performed using an E500 Elexsys Bruker spectrometer operating at 9.25-9.3 GHz, equipped with a high sensitivity continuous wave (CW) resonator. Spectra were recorded at low temperature (135 ± 5 K) at a microwave power of 20.0 mW, modulation amplitude of 5.0 G, a time constant of 120 ms, and receiver gain of 60.0 dB. The samples were measured in a 1.6-mm quartz tube (Wilmad-LabGlass, Vineland, NJ) placed in a 4.0 mm quartz tube for cooling. CW-EPR simulations were carried out using MATLAB, with the EasySpin toolbox[7].

Sample preparation

200 µL of 8 µM TasA solution was mixed with 2 µL of 100X stock solution of $CuCl_2$ to reach final concentrations of 0.1, 0.5, 1, 3, and 5 mM $CuCl_2$. Samples were loaded Capillary. Gel samples were prepared by mixing 2 mL of 8 µM TasA with 2 µL of 100X stock solution of $CuCl_2$ to reach final concentrations of 0.1, 0.5, 1, 3, and 5 mM $CuCl_2$, and 20 µL of 2M $ZnCl_2$ then centrifuged 4600xg, 5 min. Gels were placed in NMR tube and measured.

X-ray photoelectron spectroscopy (XPS)

Samples were prepared as follows, 1 mL of 12 µM TasA mixed with 10 µL of 2M ZnCl2 (final concentration is 20mM) then centrifuged 4600xg, 5 min. TasA-Zn gels were washed in triple distilled water for 16 h then lyophilized, as well as pure TasA solution. TasA, and TasA-Zn gel powders were then measured with the Axis Supra (Kratos) spectrometer.

Small angle X-ray scattering (SAXS)

TasA-Zn samples were prepared by mixing 1 mL of 12 µM TasA with 10 µL of 100X stock $ZnCl_2$ solution to reach final concentrations of 1 and 20 mM $ZnCl_2$ (similarly to cryo-TEM samples). Samples were then loaded into a 2 mm flow through cell setup of ID02 beamline, ESRF. Data were averaged and background subtracted using the SAXSutilities2 software. Data for the plots were averaged over at least 20 different points for each sample (exact numbers are indicated in figure captions).

X-ray Diffraction (XRD)

TasA-Zn hydrogel was prepared 1 mL of 20 µM TasA with 10 µL of 2 M stock $ZnCl_2$ solution to 20 mM, placed between two silicon nitride membranes that were sealed to keep the sample humid. Samples were measured at ID13 beamline of the European Synchrotron Radiation Facility (ESRF) (Grenoble), using intermediate beam size (200 nm). Data was background subtracted with Origin2024.

Fourier Transform Infrared Spectroscopy (FTIR)

A 1 mL of 12 µM TasA with 10 µL of 100X stock $ZnCl_2$ solution to reach final concentrations of 1, 3, 5,10, and 20 mM. Centrifuged 4600xg, 5 min and washed with triple distilled water for 16 h. Then froze with liquid nitrogen and lyophilized. TasA, and TasA-Zn gel powders were then measured using NICOLET iS50 FT-IR (Thermo Scientific), with SMART iTX accessory.

Atomic force microscopy (AFM)

A droplet of 10 µL of 6 µM TasA and 6 µM TasA with 1 mM $ZnCl_2$ was placed on freshly cleaved mica, incubated for 10 min then measured with The Cypher-ES AFM microscope.

#### References


[1] D. Romero, C. Aguilar, R. Losick, R. Kolter, "Amyloid fibers provide structural integrity to *Bacillus subtilis* biofilms" *Proceedings of the National Academy of Sciences* **2010**, *107*, 2230–2234.

[2] L. Chai, D. Romero, C. Kayatekin, B. Akabayov, H. Vlamakis, R. Losick, R. Kolter, "Isolation, Characterization, and Aggregation of a Structured Bacterial Matrix Precursor" *Journal of Biological Chemistry* **2013**, *288*, 17559–17568.

[3] S. S. Branda, J. E. González-Pastor, S. Ben-Yehuda, R. Losick, R. Kolter, "Fruiting body formation by *Bacillus subtilis*" *Proceedings of the National Academy of Sciences* **2001**, *98*, 11621–11626.

[4] H. Salat, R. Murcio, E. Arcaute, "Multifractal methodology" *Physica A: Statistical Mechanics and its Applications* **2017**, *473*, 467–487.

[5] A. Noa, H.-S. Kuan, V. Aschmann, V. Zaburdaev, L. Hilbert, "The hierarchical packing of euchromatin domains can be described as multiplicative cascades" *PLoS Comput. Biol.* **2021**, *17*, e1008974.

[6] H. Atmanspacher, H. Scheingraber, G. Wiedenmann, "Determination of f(α) for a limited random point set" *Phys. Rev. A (Coll Park).* **1989**, *40*, 3954–3963.

[7] S. Stoll, A. Schweiger, “EasySpin, a comprehensive software package for spectral simulation and analysis in EPR” *Journal of Magnetic Resonance* **2006**, *178*, 42–55.
